# Defying Salinity, Drought, and pH Extremes: A Multifunctional Rhizobacterium, *Burkholderia gladioli* ST3M-39a, Matches Fertilizer Efficacy in Wheat via Phosphate Solubilization

**DOI:** 10.1101/2025.09.01.673583

**Authors:** Sudip Silwal, Asmita Shrestha, Shreejan Pokharel, Bignya Chandra Khanal, Ramesh Acharya, Gyanu Raj Pandey

**Author notes:** Corresponding Author: (GRP). These authors contributed equally to this work (Sudip Silwal, Asmita Shrestha).

## Abstract

Global phosphorus scarcity and the environmental impacts of chemical fertilizers necessitate sustainable microbial alternatives for agriculture. We characterized *Burkholderia gladioli* ST3M-39a, a maize rhizosphere isolate, as a multifunctional plant growth-promoting rhizobacterium with exceptional climate resilience. The strain achieved rapid phosphate solubilization (177.96 ± 5.26 µg/mL within 24 h; molybdenum-antimony assay), zinc solubilization, and ammonia production, EPS production, and produced stress-alleviating enzymes (cellulase and protease). Crucially, it maintained robust growth and phosphate-mobilizing capacity under extreme abiotic stresses: pH 4.5-8.5, 7.5% NaCl salinity, and drought-mimicking low water activity (a_w_ 0.950, 32% sorbitol). In wheat trials, ST3M-39a inoculation significantly increased the growth parameters (p < 0.05 vs. those of the uninoculated controls), resulting in 85-92% of the biomass stimulation observed with diammonium phosphate (DAP) fertilizer. This multifunctional stress tolerance, coupled with its near-fertilizer efficacy, positioned ST3M-39a as a transformative bioinoculant for degraded soils. Field validation of its agricultural deployment and ecological impact is now pivotal.

## 1 Introduction

Establishing a sustainable agricultural system is a paramount global challenge driven by rising food demands. Meeting these demands through increased crop yields risks indiscriminate agrichemical use, causing environmental degradation (1). In addition to resource depletion, biodiversity loss, and climate change, several abiotic stresses, including salinity, drought, cold, and metal toxicity, severely impair crop growth, development, and productivity worldwide (2). An alternative strategy leverages beneficial plant microbiomes, known as plant growth-promoting bacteria (PGPB). These bacteria help plants acquire essential nutrients, promote growth, modulate phytohormones, and increase stress tolerance, offering an economically viable solution with significant agronomic utility (3,4).

Phosphorus is an indispensable plant macronutrient essential for nucleic acid synthesis, cell division, and critical cellular processes, including photosynthesis, carbohydrate metabolism, energy production, redox homeostasis, and signal transduction (5–7). Its deficiency remains a primary growth-limiting factor in terrestrial ecosystems, as soil solution concentrations of orthophosphate (the plant-available form) rarely exceed 10 µM despite substantial total soil phosphorus reserves (8). Crucially, soil phosphorus pools are dominated by recalcitrant organic compounds (30–65% of total P) and inorganic phosphorus (35-70%) that rapidly precipitate with metal cations (Fe³⁺, Al³⁺, Ca²^+^). This results in fertilizer-derived phosphorus accumulating primarily in plant-unavailable forms, sustaining phosphorus limitation in agricultural soils despite repeated fertilization and consequently reducing agricultural productivity (9), (10,11).

The widespread application of phosphate fertilizers results in low recovery efficiency (10-30%), with residual phosphorus either becoming fixed in soils or contaminating aquatic systems through runoff. This accelerates eutrophication and expands agriculture’s carbon footprint (12),(13). The accumulation of soil phosphates is estimated to be sufficient to sustain global crop production for at least a century. Despite these reserves, chemical fertilizer production and use continue unabated worldwide (14). Economic projections indicate that commercially viable phosphate reserves may be depleted within 100-200 years at current extraction rates (15). Fertilizer prices surged 292.7% between 2000-2022 due to crop production inflation and rising global food costs. Furthermore, mining and processing phosphate fertilizers, which contain natural radioactive elements, compromise public health (14). Given the environmental damage from fertilizer overuse, underutilization of legacy phosphorus, and dwindling global reserves, developing sustainable strategies to enhance soil phosphorus bioavailability is imperative for food security (5).

Phosphate-solubilizing bacteria (PSB) offer a sustainable strategy to mitigate soil phosphorus deficiency by enhancing bioavailable phosphorus through three key mechanisms: (1) acid-mediated mineralization of fixed phosphorus pools, (2) enzymatic hydrolysis of organic compounds, and (3) chelation of metal ions (Fe³⁺, Al³⁺, Ca²^+^) via extracellular polysaccharides and organic acids. These processes collectively reduce the soil pH while liberating orthophosphates (16). As biofertilizers, PSB enhance plant growth, nutritional status, root architecture, stress tolerance, and competitive ability (17). Documented growth improvements in maize, wheat, paddy, rice, pea, and lettuce confirm their efficacy across diverse crops (18–21), with specific studies demonstrating significant increases in wheat biomass and phosphorus content following PSB inoculation (17). In addition to phosphorus solubilization, these microorganisms function as phytohormone producers, biocontrol agents against pathogens, heavy-metal bioremediation agents, and abiotic stress mitigators (22).

Bacterial genera including *Pseudomonas*, *Bacillus*, *Rhizobium*, *Burkholderia*, *Azotobacter*, *Microbacterium*, and *Azospirillum* represent the most significant PSB (23,24). In low-phosphorus soil, *Burkholderia* species enhance maize biomass by 15-30% and increase bioavailable phosphorus in the rhizosphere by 20-50% (25). As high-performance PSB, *Burkholderia* species exhibit broad biotechnological potential, promoting plant growth and development while conferring resistance against biotic and abiotic stresses (26). Notably, *B. gladioli* have robust plant growth-promoting effects through multiple mechanisms: phosphate solubilization, nitrogen fixation, indole-3-acetic acid (IAA) production, ACC (1-aminocyclopropane-1-carboxylate) deaminase activity, siderophore secretion, and phytohormone synthesis. These traits stem from its capacity to biosynthesize diverse bioactive secondary metabolites, positioning *B. gladioli* as a prime candidate for ecological fertilizer development (27–31).

This study isolated and characterized *Burkholderia gladioli* ST3M-39a from maize rhizosphere soil as a multifunctional plant growth-promoting rhizobacterium (PGPR) with high phosphate-solubilization activity. We systematically evaluated its: (i) phosphate solubilization efficiency under extreme abiotic stresses, (ii) other plant growth promoting traits, and (iii) functional impact on wheat growth in greenhouse trials, demonstrating near-fertilizer efficacy.

## 2 Materials and methods

### 2.1 Materials and equipment

All microbiological growth media and general reagents were procured from HiMedia and Fisher Scientific. Molecular biology-grade reagents, buffers, and PCR master mix were sourced from Amresco and New England Biolabs, respectively, with PCR primers synthesized by Macrogen (South Korea). Laboratory glassware was supplied by Borosil. Microbial characterization employed an Olympus CX21i microscope. DNA extraction and other centrifugation utilized a REMI NEYA 16R centrifuge, PCR amplification was performed on a MiniAmp^TM^ Plus Thermal Cycler (Thermo Fisher Scientific), and spectrophotometric measurements used a Labdex LX111VS instrument.

### 2.2 Isolation and maintenance of organisms

Organisms were isolated from rhizospheric soil of maize plants collected in Dumre beshi, Bharatpur-29, Chitwan, Nepal (27°48’52’’N and 084°28’47’’E) as shown in Fig 1. The soil samples (1 g) were homogenized in 10 ml of sterile water, serially diluted, and spread onto yeast mannitol agar. The plates were incubated aerobically at 30℃ for 24 h. Morphologically distinct bacterial colonies were subcultured on glucose-yeast extract-peptone (GYP; 1% glucose, 0.5% yeast extract, 0.3% peptone) agar to obtain pure cultures. Isolates were preserved as glycerol stocks at -80℃ for long-term storage and maintained on bacterial slants (stored at 4℃) for experimental use (32).

**Figure 1:**
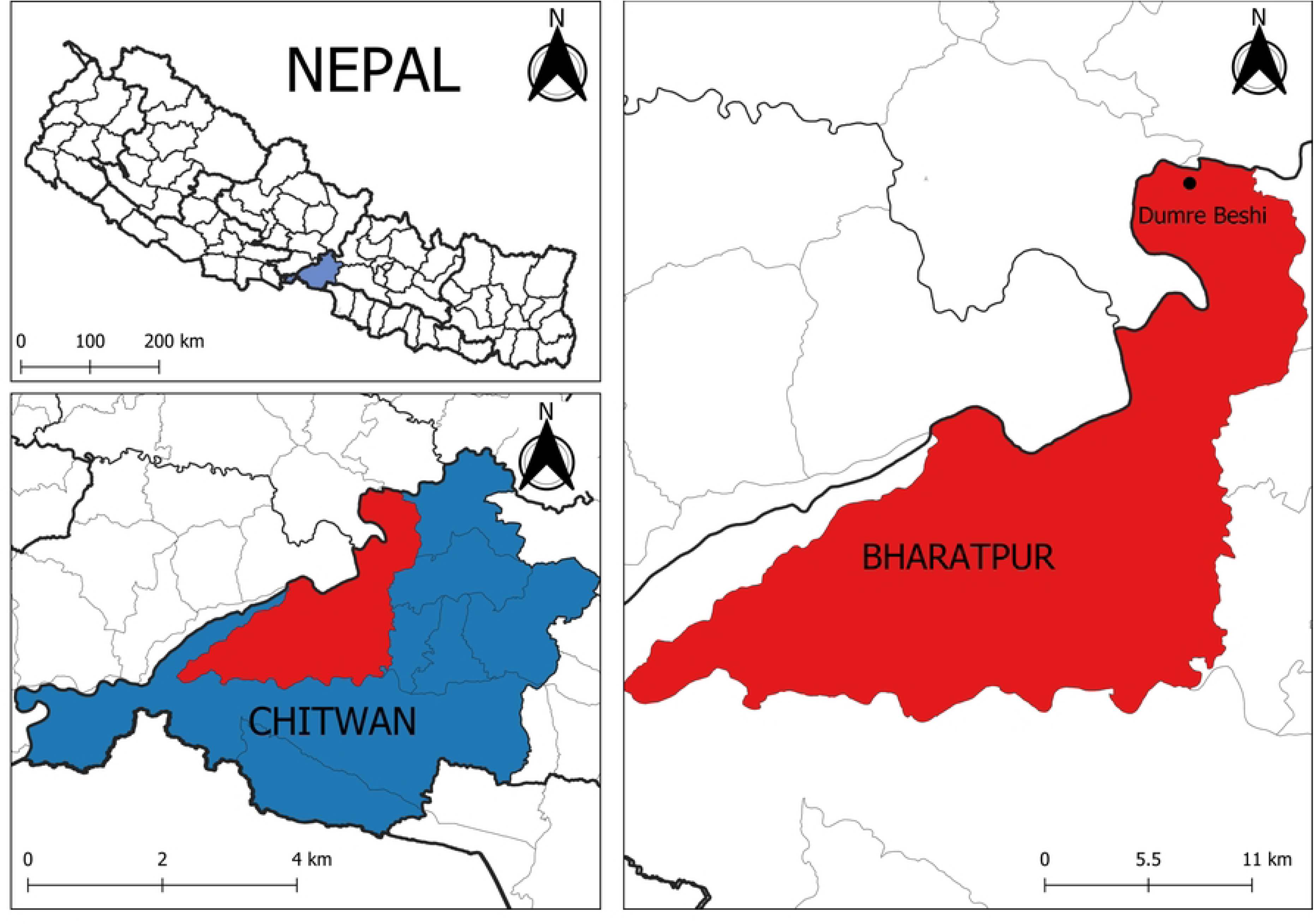
Location of the sampling site: Dumre beshi, Bharatpur 29, Chitwan, Nepal

### 2.3 Screening for phosphate solubilization

#### 2.3.1 Qualitative assay

Isolates were screened for phosphate solubilization on Pikovskaya’s agar (PKA) (33) and National Botanical Research Institute’s phosphate (NBRIP) media (34). Both media were incubated at 30℃ for 72 h under aerobic conditions. Colonies that formed halo zones on both media were identified as phosphate solubilizers. Isolate ST3M-39a, the sole isolate exhibiting phosphate solubilization on both media, was selected for further analysis. The phosphate solubilization index (PSI) was calculated as follows:

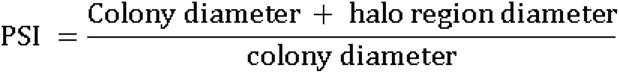

#### 2.3.2 Quantitative assay

The strain exhibiting the sole halo zone was selected for quantitative phosphate solubilization analysis. The strain was precultured in nutrient broth (24 h, 30°C, 100 rpm) before the standardized inoculum (OD_600_ 0.6; 2-10% v/v) was aseptically transferred to 30 mL NBRIP medium (pH 6.8) in 250 mL flasks, with uninoculated medium used as a negative control. Cultures were incubated aerobically (30℃, 100 rpm), and 1 mL samples were collected after 24 h for phosphate quantification. After centrifugation (10,000 × g, 5 min, 4℃), the supernatant was analyzed for solubilized phosphate via the molybdenum-antimony colorimetric method (35). The final pH of the medium was also recorded. All the experiments were performed in triplicate.

### 2.4 Identification of organism

#### 2.4.1 Molecular identification

##### 2.4.1.1 DNA extraction and sequencing

The bacterial isolate was incubated in nutrient broth at 30°C for 24 h. Genomic DNA was extracted via the Sambrook and Russell method (36), and the 16S rRNA gene was amplified with the universal primers 27F (5’-AGAGTTTGATCMTGGCTCAG-3’) and 1492R (5- TACGGYTACCTTGTTACGACTT-3’). Sequencing was performed by Macrogen (South Korea). The raw sequences were assembled and trimmed in a codon code aligner, and the consensus sequence was subsequently deposited in NCBI GenBank (accession OR143748.1). For phylogenetic analysis, the sequence was compared to the NCBI database via BLASTN (16S rRNA dataset), and 20 closely related sequences were retrieved in FASTA format.

##### 2.4.1.2 Maximum likelihood analysis of taxa

A maximum likelihood phylogenetic tree was constructed in MEGA12 (37) via the Tamura-Nei model (38), with nodal support assessed via 2,000 bootstrap replicates (39).

**1. Morphological and biochemical characterization:**

The isolate was identified morphologically and biochemically following *Bergey’s Manual of Systematic Bacteriology* (40).

### 2.5 Growth under different abiotic conditions

Bacterial growth was assessed in nutrient broth under controlled abiotic stresses: pH gradients (4.0-11.5; 0.5-1.0 increments), NaCl concentrations (0-9.0% w/v; pH 6.8), and sorbitol-induced osmotic stress (0-40% w/v; pH 6.8). Water activity (a_w_) for sorbitol solutions was calculated following established methods (41). Overnight cultures (OD_600_ = 0.6) were aseptically inoculated at 2% (v/v) into fresh NBRIP medium supplemented with test conditions. Cultures underwent aerobic incubation at 30°C with 100 rpm agitation for 48 h. All treatments were replicated in triplicate.

### 2.6 Phosphate solubilization under different abiotic conditions

Phosphate solubilization was assessed in NBRIP medium under variable stress conditions: pHs (4.5, 7, and 8.5), NaCl (0-7.5% w/v, 2.5% increments; initial pH 6.8), and sorbitol-induced osmotic stress (0-32% w/v, 8% increments; initial pH 6.8). Overnight nutrient broth culture (OD_600_ = 0.6) was inoculated at 6% (v/v) into test media. Following 24 h aerobic incubation at 30°C with 100 rpm agitation, solubilized phosphate was quantified using previously described method and final pH recorded. All assays were performed in triplicate.

### 2.7 Other PGPR properties

#### 2.7.1 Zinc solubilization

Zinc solubilization was assessed by culturing the isolate in mineral salt media supplemented with 0.1% zinc oxide (20). The plates were incubated at 30℃ for 7 days, and the zinc solubilization efficiency (ZSE%) was calculated as (clearing zone diameter/colony diameter) × 100 (42).

#### 2.7.2 Exopolysaccharide production

Exopolysaccharide (EPS) production was induced by inoculating 100 mL LB broth with 100 μL bacterial culture (OD_600_ = 0.6) followed by 96 h incubation at 30°C with 100 rpm orbital shaking. Cells were pelleted by centrifugation, and the supernatant treated with two volumes of ice-cold 95% ethanol to precipitate EPS. Precipitates were recovered, dried, and quantified gravimetrically (43,44).

#### 2.7.3 Cellulase activity

For cellulolytic activity, the isolates were grown in basal salt media supplemented with 1% carboxymethyl cellulose (30°C, 48 h). After incubation, the agar plate was stained with 0.1% Congo red solution for 15 min, followed by counterstaining with 1 M NaCl (20 min). Cellulose hydrolysis was identified by clear zones surrounding colonies (45).

#### 2.7.4 Protease activity

For the protease test, the isolate was cultured on skim milk powder agar (5% skim milk powder) and incubated at 30℃ for 48 hours. The clear halo region indicates the production of protease by the isolate (46).

#### 2.7.5 Ammonia production

Ammonia production was qualitatively evaluated via the method by Chen *et al*. (47) by inoculating test strains in peptone broth and incubating them at 30°C for 48 h. Following incubation, 500 μL of Nessler’s reagent was added, with the development of a yellow to brown color indicating presence of ammonia.

### 2.8 Greenhouse experiment

A controlled pot trial was conducted using 45 sterile 9-inch diameter pots filled with autoclaved soil. The wheat (*Triticum aestivum*) was sown, and germination occurred one-week post-sowing. The seedlings were thinned to five uniform plantlets per pot and maintained under greenhouse conditions. The treatments included: (A) a negative control (no supplements; 15 pots), (B) a positive control (90 kg/ha diammonium phosphate [DAP]; 15 pots), and (C) test samples (1 mL of phosphate-solubilizing bacterial inoculum, 1 × 10^9^ CFU/mL; 15 pots). Pots received 50 mL autoclaved water weekly.

At maturity, agronomic parameters shoot length, root length, spike length, chlorophyll content (SPAD meter), dry biomass, yield per pot, and seed number per pot were measured (17).

### 2.9 Statistical analysis

The data were analyzed via Origin Pro and Microsoft Excel. Quantitative variables were assessed by ANOVA, with Tukey’s test employed for pairwise mean comparisons at a 95% confidence interval.

## 3 Results

### 3.1 Screening for phosphate solubilization

The tested isolate exhibited significant phosphate-solubilizing activity, as evidenced by halo zone formation on both Pikovskaya’s (PVK) and National Botanical Research Institute’s Phosphate (NBRIP) agar media (Fig 2I). The phosphate solubilization index (PSI) was comparable in both media, with values of 1.27 ± 0.097 (PVK) and 1.41 ± 0.10 (NBRIP), indicating efficient mineral phosphate solubilization. Further quantitative analysis revealed that the inoculum size significantly influenced the phosphate solubilization efficiency (Fig 2II). The highest soluble phosphate concentration (148.97 ± 7.45 µg/mL) was achieved with the 10% inoculum, which was significantly greater (p < 0.05) than that obtained with the 4% (101.52 ± 4.13 µg/mL) and 2% (89.58 ± 4.93 µg/mL) inoculum sizes. However, since the phosphate solubilization ability of 6% (129.65 ± 2.97 µg/mL) and 8% (144.71 ± 3.98 µg/mL) inoculum was not significantly different from that of the 10% inoculum, the 6% inoculum was selected for subsequent experiments. The initial pH of the medium (pH 6.8) decreased after the incubation period. The most pronounced reduction (pH 5.02 ± 0.01) occurred with the 10% inoculum. The pH decline became less pronounced with decreasing inoculum size: 8% (pH 5.07 ± 0.01), 6% (pH 5.09 ± 0.02), 4% (pH 5.54 ± 0.03), and 2% (pH 5.62 ± 0.02) as shown in Figure 2II. However, the pH decline was similar for the 6%, 8%, and 10% inoculum sizes.

**Figure 2:**
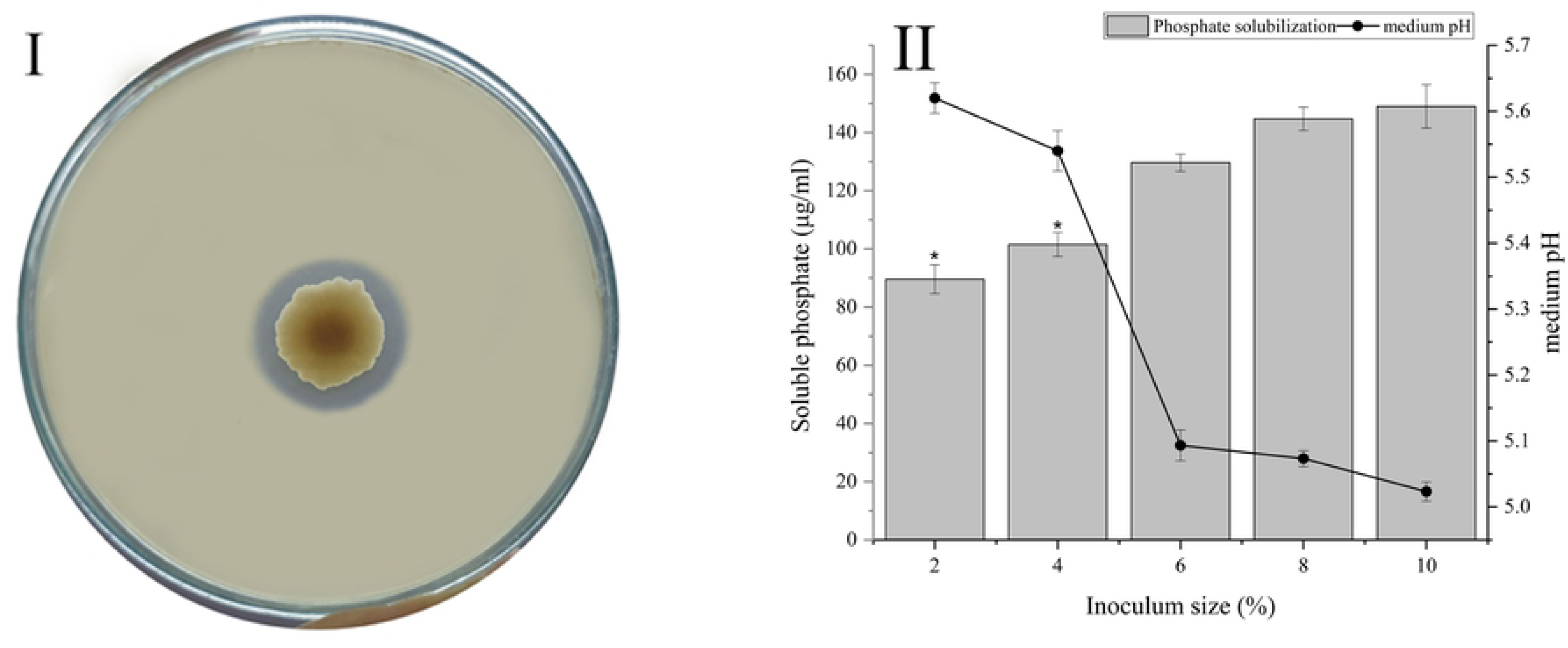
(I) Halo zone formation by the isolate ST3M-39a on Pikovskaya’s (PVK) agar. (II) Soluble phosphate released and final medium pH in NBRIP medium across inoculum doses. *Significantly lower than 10% inoculum (α = 0.05). Data represent mean ± SE (n = 3).

### 3.2 Identification of the organism

#### 3.2.1 Molecular characterization

Comparisons using nucleotide BLAST of the 16S rRNA sequence of strain ST3M-39a revealed 99.81% identity with *Burkholderia gladioli* strain NBRC 13700, 99.53% identity with *Burkholderia plantarii* strain CIP 105769, 99.41% identity with *Burkholderia perseverans* strain INN12, and 99.16% identity with *Burkholderia glumae* strain P 1-22-1. Maximum likelihood analysis revealed that strain ST3M-39a clustered robustly within the *B. gladioli* monophyletic clade, exhibiting the closest phylogenetic affinity to reference strains of *B. gladioli*, with all strains forming a highly supported subclade (bootstrap > 95%) distinct from other *Burkholderia* species, including the phytopathogens *B. glumae* (LMG 2196) and *B. plantarii* (NIAES 1723), opportunistic pathogen *B. cenocepacia* (LMG 16656), and environmental species *B. stagnalis* (LMG 28156), as shown in Fig 3.

**Figure 3:**
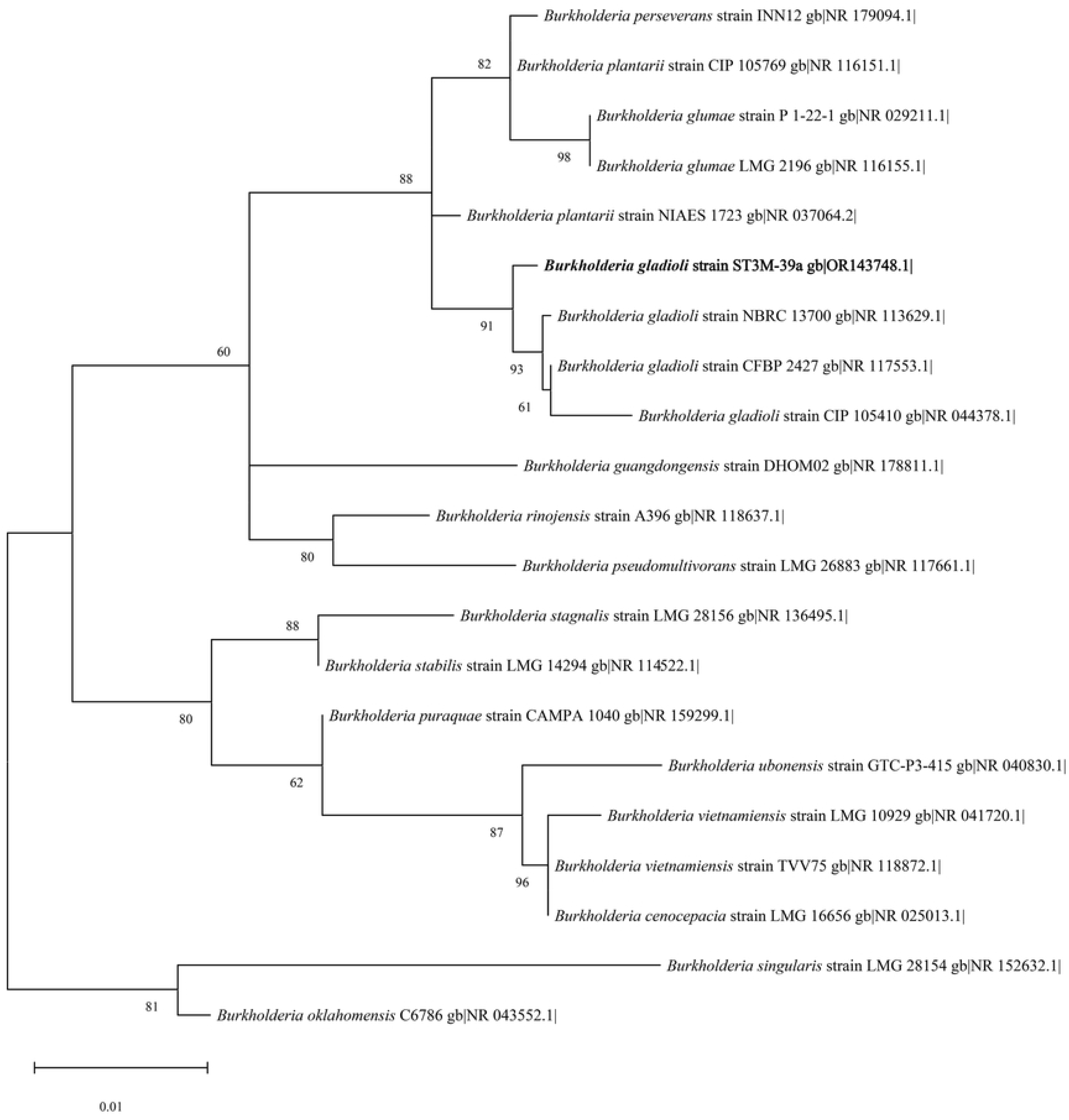
Maximum likelihood phylogenetic tree of the isolate based on 16S rRNA sequence data (Tamura-Nei model. Bootstrap values (2000 replicates) are shown at nodes; branches with <50% support were collapsed. Initial topologies were generated via neighbor-joining and BioNJ algorithms, with the optimal tree selected by log-likelihood.

#### 3.2.2 Morphological and biochemical characterization

### 3.3 Growth under different abiotic conditions

#### 3.3.1 pH levels

The isolate ST3M-39a grew over a wide pH range (5–11) (Fig 4I). The growth at neutral pH (2.023 ± 0.059) was compared with that at acidic and basic pH. Growth was significantly lower at pH values of 4 (0.0053 ± 0.0014), 4.5 (0.048 ± 0.021), 5 (0.28 ± 0.0045), 10 (1.782 ± 0.047), 11 (0.307 ± 0.041), and 11.5 (0.052 ± 0.0032) than at pH 7 (p < 0.05). However, no significant differences were observed between pH 7 and pH 5.5 (2.108 ± 0.426), 6 (2.027 ± 0.078), 8 (1.997 ± 0.024), or 9 (2.092 ± 0.043) (p < 0.05). So, the optimal pH range for the isolate was 5.5-9.

**Figure 4:**
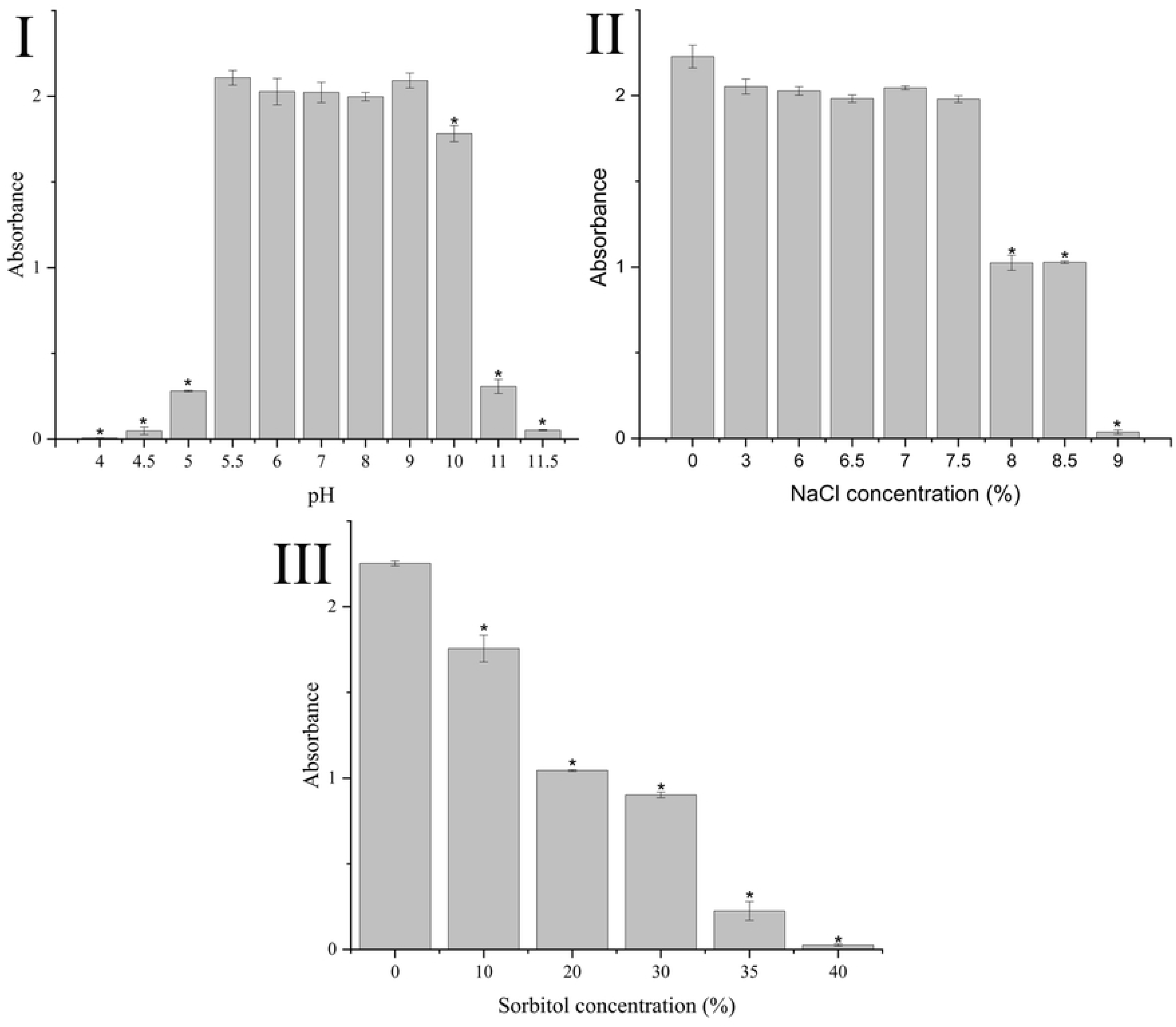
(I) Growth of ST3M-39a in nutrient broth at various pH values. *Significantly lower than that at pH 7 (α = 0.05). (II) Growth of ST3M-39a under NaCl gradients. *Significantly lower than growth at 0% salt concentration (α = 0.05). (III) Growth of ST3M-39a under sorbitol-induced osmotic stress. *Significantly lower than growth at 0% sorbitol concentration (α = 0.05). The data represent means ± SEs (n = 3).

#### 3.3.2 NaCl concentrations

The isolate exhibited growth across a broad NaCl tolerance range (0-8.5%), with absorbance values ranging from 1.027 to 2.228 (Fig 4II). While optimal growth occurred without NaCl (2.228 ± 0.066), significant growth reduction was observed at higher concentrations of 8% (1.024 ± 0.0438), 8.5% (1.027 ± 0.007), and 9% (0.037 ± 0.012) (p < 0.05). Growth remained comparable to that of 0% NaCl at concentrations (ranging from 3-7.5%), with no significant differences observed at 3% (2.053 ± 0.044), 6% (2.028 ± 0.025), 6.5% (1.982 ± 0.022), 7% (2.045 ± 0.012), and 7.5% (1.980 ± 0.020) NaCl concentrations (p > 0.05). Therefore, the isolate exhibited optimal growth at a salt concentration range of 0-7.5%.

#### 3.3.3 Osmotic stress

The isolate grew over a broad range of sorbitol concentrations (0%-35%), with absorbance values ranging from 0.225 to 2.253 as shown in Fig 4III. Optimal growth was detected in the absence of sorbitol (2.253 ± 0.014), with significantly reduced growth observed at higher concentrations of 10% (1.756 ± 0.078; a_w_ = 0.989), 20% (1.044 ± 0.005; a_w_ = 0.974), 30% (0.902 ± 0.016; a_w_ = 0.955), 35% (0.225 ± 0.055; a_w_ = 0.943), and 40% (0.026 ± 0.006; a_w_ = 0.929) (p < 0.05).

### 3.4 Phosphate solubilization under different conditions

#### 3.4.1 pH levels

The isolate demonstrated significant phosphate solubilization capacity across various pH conditions (4.5, 7.0, and 8.5), with soluble phosphate concentrations ranging from 108.42 to 133.34 µg/mL (Fig 5I). Quantitative analysis revealed pH-dependent solubilization efficiency, with maximum phosphate release observed at neutral pH (133.35 ± 3.23 µg/mL at pH 7.0). Solubilization was significantly reduced under both acidic and alkaline conditions, yielding 118.29 ± 1.73 µg/mL at pH 4.5 and 108.42 ± 1.30 µg/mL at pH 8.5 (p < 0.05 for both comparisons versus pH 7.0). Following incubation, the final medium pH was 5.21 ± 0.01 (initial pH of 4.5), 5.03 ± 0.01 (initial pH of 7), and 5.32 ± 0.01 (initial pH of 8.5). The most significant pH decrease occurred in the culture initially adjusted to a neutral pH (7.0) as shown in Fig 5I.

**Figure 5:**
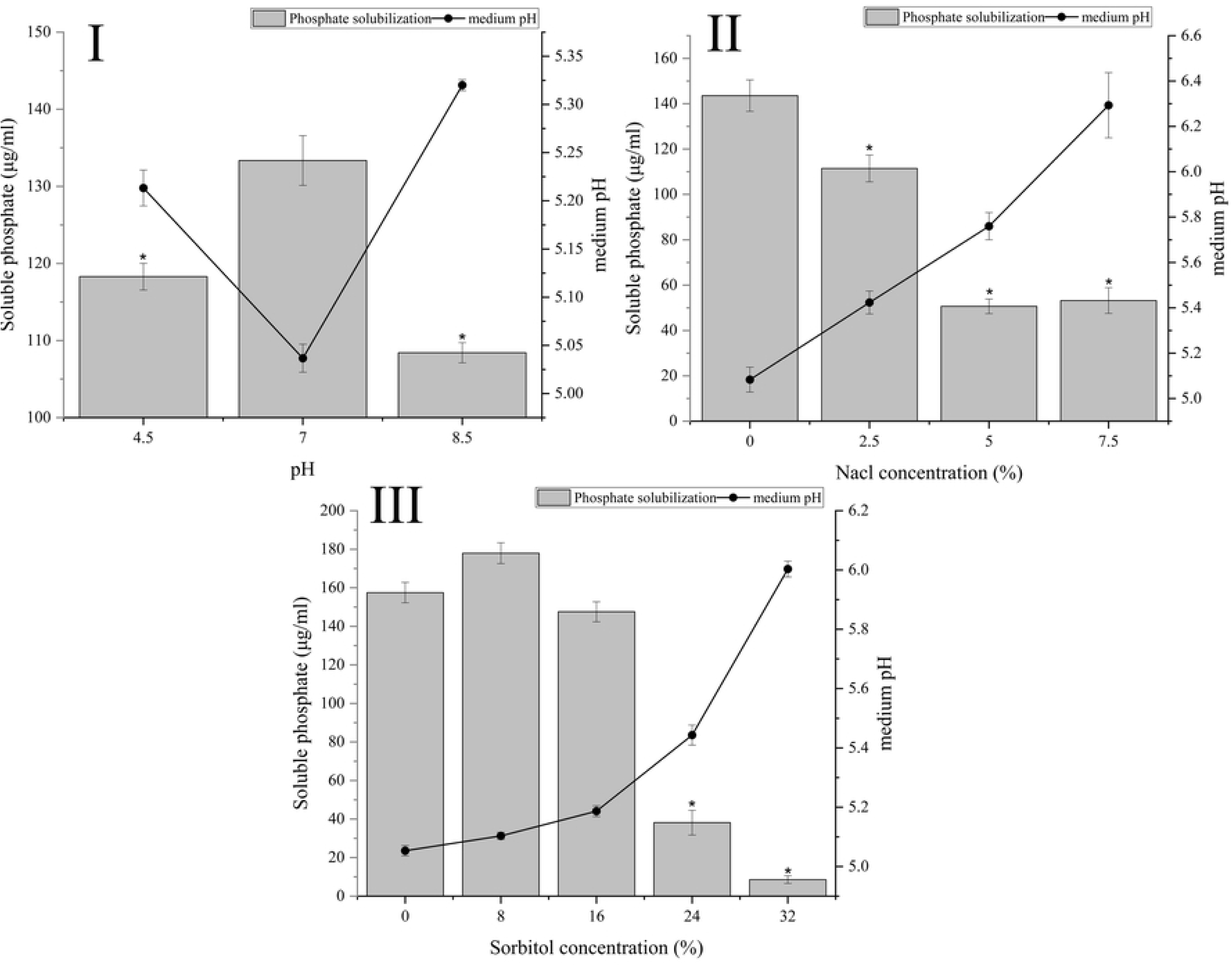
(I) Soluble phosphate released and final medium pH in NBRIP medium under a pH gradient. *Significantly lower than that at pH 7 (α = 0.05). (II) Soluble phosphate release and final medium pH in NBRIP medium under NaCl-induced salinity. *Significantly lower than the 0% NaCl concentration (α = 0.05). (III) Soluble phosphate release and final medium pH in NBRIP medium under sorbitol-induced osmotic stress. *Significantly lower than 0% sorbitol concentration (α = 0.05). The data represent means ± SEs (n = 3).

#### 3.4.2 NaCl concentration

The isolate exhibited different phosphate solubilization capacities across a range of NaCl concentrations (0-7.5%), releasing soluble phosphate concentrations ranging from 50.66 to 143.58 µg/mL (Fig 5II). The maximum solubilization occurred in the absence of NaCl (143.58 ± 6.94 µg/mL). This activity was significantly inhibited by increasing salinity, with phosphate concentrations decreasing to 111.47 ± 5.91 µg/mL at 2.5% NaCl and further decreasing to 50.66 ± 3.23 µg/mL at 5% NaCl and 53.21 ± 5.68 µg/mL at 7.5% NaCl (p < 0.05 for all comparisons versus the 0% NaCl control). Similarly, the initial medium pH of 6.8 decreased during phosphate solubilization across all NaCl concentrations (Fig 5II). The most pronounced reduction occurred without NaCl (final pH 5.08 ± 0.05), with progressively smaller decreases observed at higher salt concentrations: pH 5.4 ± 0.05 (2.5% NaCl), pH 5.76 ± 0.06 (5% NaCl), and pH 6.29 ± 0.14 (7.5% NaCl).

#### 3.4.3 Osmotic stress

The isolate exhibited varying phosphate solubilization capacities across different sorbitol concentrations (0-32%), releasing soluble phosphate concentrations ranging from 8.59 to 177.96 µg/mL (Fig 5III). Although solubilization at 8% (177.96 ± 5.26 µg/mL; a_w_ = 0.991) and 16% sorbitol (147.55 ± 5.17 µg/mL; a_w_ = 0.981) remained comparable to that of 0% control (157.50 ± 5.26 µg/mL; a_w_ = 1.0; p > 0.05), a significant reduction occurred at higher concentrations, decreasing to 38.15 ± 6.42 µg/mL at 24% (a_w_ = 0.967) and 8.59 ± 1.99 µg/mL at 32% (a_w_ = 0.950) sorbitol (p < 0.05 versus the control). Similarly, the initial medium pH of 6.8 decreased during phosphate solubilization at all sorbitol concentrations (Fig 5III). Maximum acidification occurred without sorbitol (5.05 ± 0.02 at 0%), with progressively smaller pH reductions observed at increasing concentrations: pH 5.1 ± 0.01 (8%), pH 5.18 ± 0.02 (16%), pH 5.44 ± 0.03 (24%), and pH 6.00 ± 0.02 (32%).

### 3.5 Other PGPR properties

#### 3.5.1 Zinc solubilization, EPS production, cellulase activity, protease activity, and ammonia production

The isolate demonstrated a zinc solubilization ability, forming a clear halo zone on mineral salt media supplemented with zinc oxide (Fig 6I), with a solubilization efficiency of 152.19% ± 7.25%. Exopolysaccharide (EPS) production was 183.33 ± 14.53 µg/mL. Cellulase production was evident through the formation of a hydrolysis zone (Fig 6II), whereas protease activity was confirmed by the presence of a clearing zone (Fig 6III). Additionally, ammonia production was verified by a distinct color change from yellow to brown in the medium broth (Fig 6IV).

**Figure 6:**
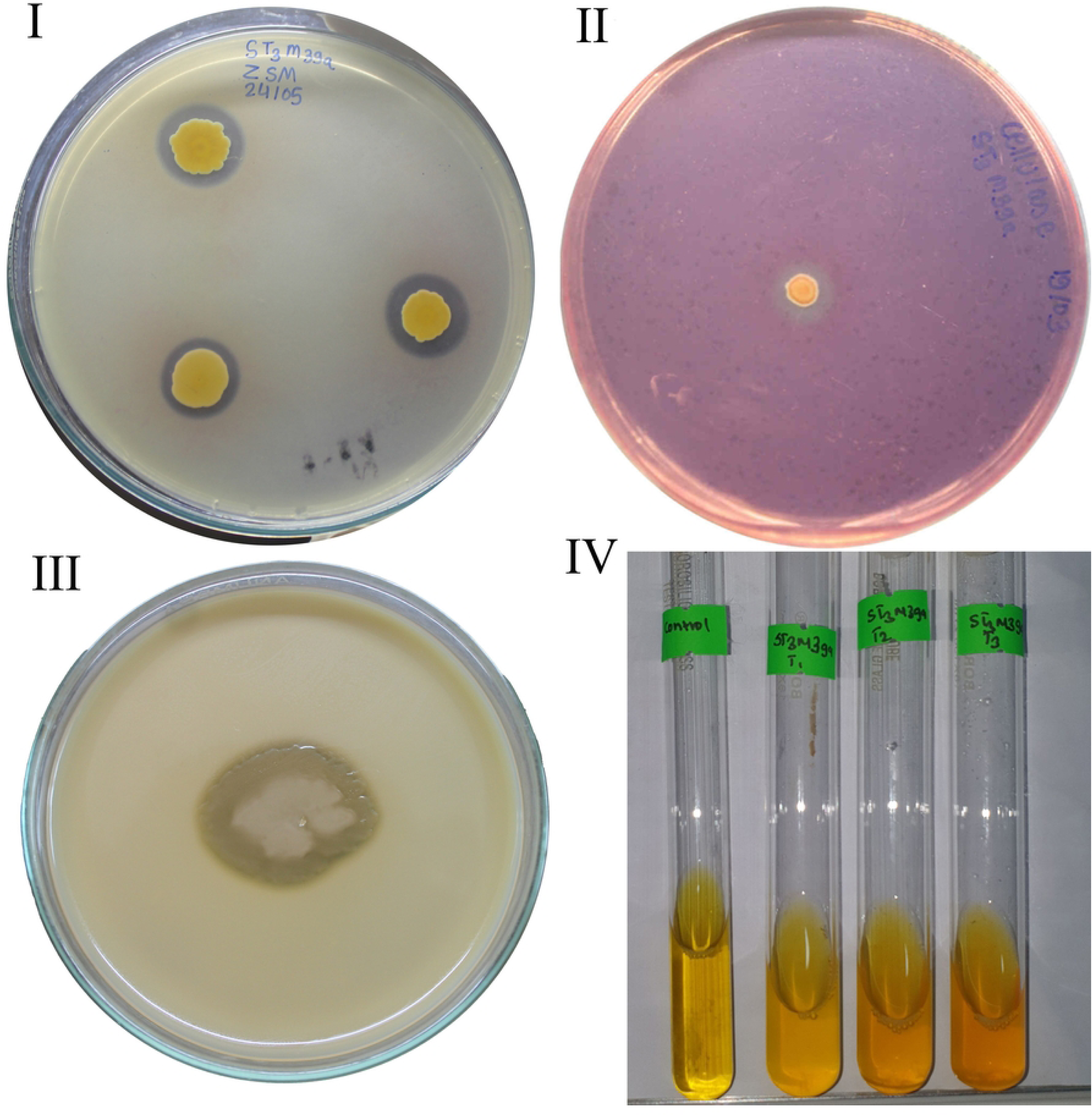
(I) Zinc solubilization by ST3M-39a on ZnO-supplemented mineral salt media (halo zone). (II) Cellulase activity on CMC agar (halo zone). (III) Protease activity on skim milk agar (halo zone). (IV) Ammonia production.

### 3.6 Greenhouse experiment

As shown in the radar plot (Fig 7), compared with the negative control samples, the test samples presented notable advantages in terms of aerial growth characteristics and yield, along with improved root development. On the other hand, compared with the test samples, the positive control samples presented enhanced growth performance with superior results in terms of chlorophyll content, root length, spike development, biomass accumulation, and yield. However, the test sample presented greater shoot length than did the positive control. Compared with negative control, both the test samples and the positive control significantly improved growth across all the measured parameters.

**Figure 7:**
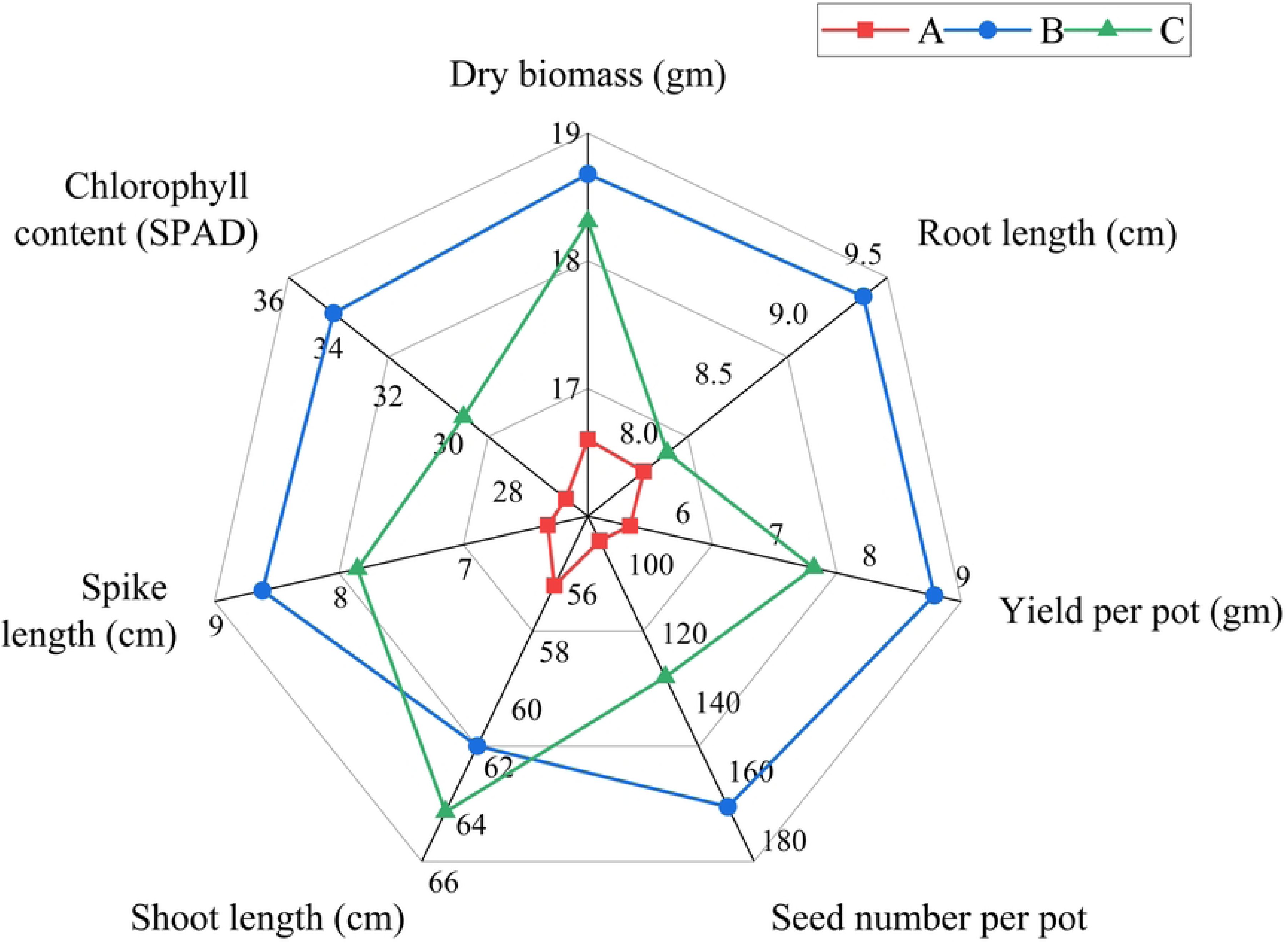
Radar plot of greenhouse trial outcomes: (A): negative control, (B): DAP-supplemented positive control, (C): ST3M-39a inoculated test samples.

#### 3.6.1 Effects on yield

##### 3.6.1.1 Chlorophyll content

Chlorophyll content analysis revealed significant variations among the treatment groups (Fig 8I). The positive control presented the highest mean chlorophyll level (34.49 ± 0.57 SPAD), representing a 14.4% increase over that of the test samples (30.15 ± 0.40 SPAD) and 29.1% greater than the negative control (26.73 ± 0.67 SPAD). The test samples maintained intermediate values, with a 12.8% higher chlorophyll content than that of negative control. Statistical analysis confirmed that these differences were significant (p < 0.05), with the positive control significantly outperforming both other treatments, whereas the test samples remained significantly superior to the negative control.

**Figure 8:**
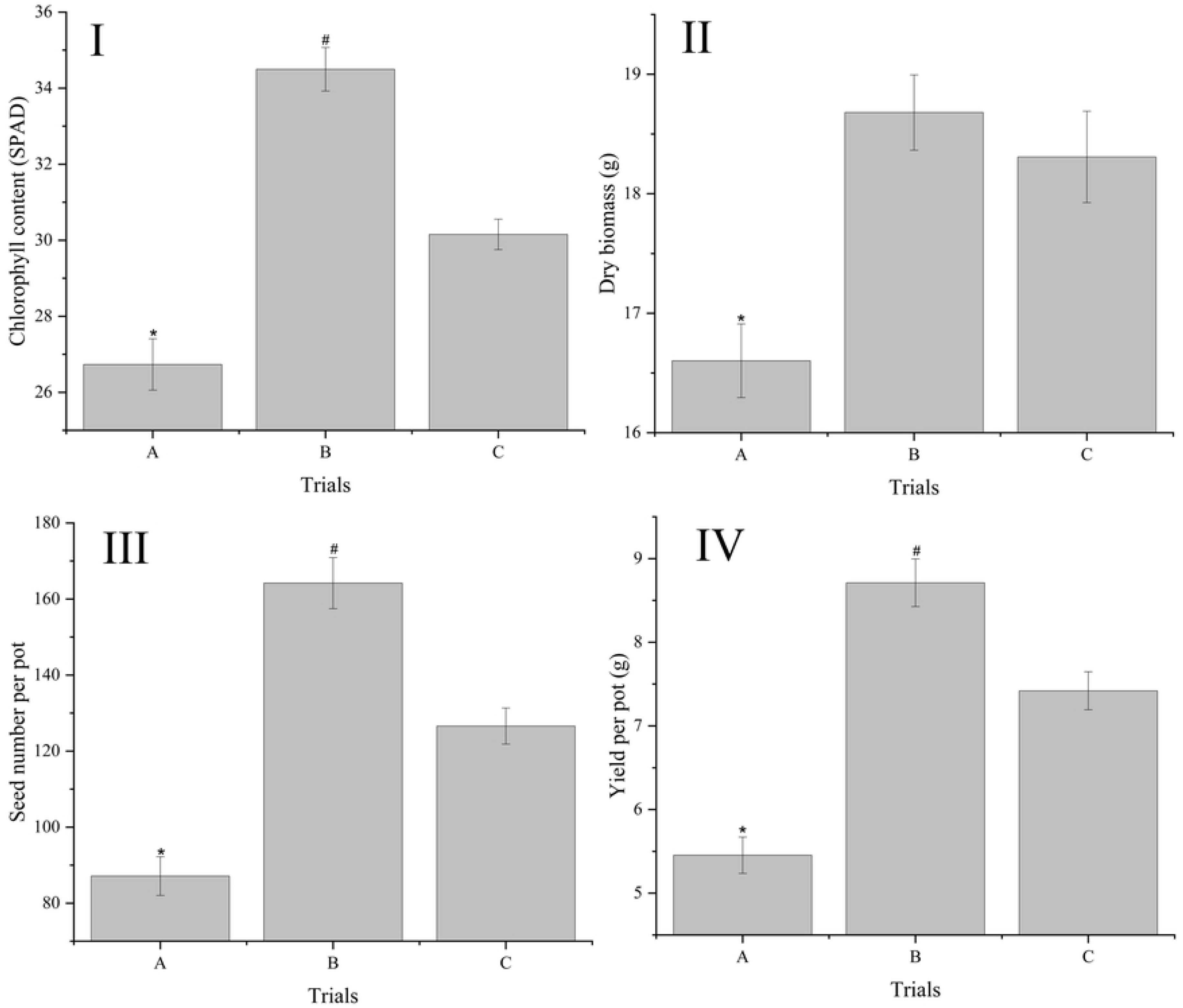
Yield parameters: (I) leaf chlorophyll content, (II) dry biomass, (III) seed count, and (IV) yield per pot. (A): Negative control, (B): DAP-supplemented positive control, (C): ST3M-39a inoculated test samples. *Significantly lower than that in trial C (α = 0.05). ^#^Significantly higher than that in trial C (α = 0.05). The data represent means ± SEs (n = 15).

##### 3.6.1.2 Dry biomass

The dry biomass of wheat varied significantly across treatments, ranging from 16.60 to 18.68 gm (Fig 8II). The test samples presented a mean dry biomass of 18.30 ± 0.38 gm, representing a significant 10.2% increase over that of negative control (16.60 ± 0.31 gm; p < 0.05). While the positive control (18.68 ± 0.31 gm) resulted in marginally greater biomass (2.1% greater than that of the test samples), this difference was not statistically significant (p < 0.05).

##### 3.6.1.3 Seed number per pot

Seed production in the wheat plants significantly varied across the treatments (Fig 8III). The test samples yielded substantially more seeds (126.6 ± 4.73 seeds) than did the negative control (87.13 ± 5.08 seeds; p < 0.05), representing a 45.3% increase. However, the number of seeds remained significantly lower than that of the positive control (164.2 ± 6.73 seeds; p < 0.05), with test samples producing 77.1% of the positive control yield.

##### 3.6.1.4 Yield per pot

Yield per pot varied significantly among treatments (Fig 8IV), with test samples (7.42 ± 0.22 g) demonstrating intermediate performance between the positive (8.71 ± 0.28 g) and negative controls (5.45 ± 0.22 g) (p < 0.05 for all pairwise comparisons). While the test samples produced 36.1% greater yield than the negative control, they achieved only 85.2% of the positive control’s yield potential.

#### 3.6.2 Effects on plant growth

##### 3.6.2.1 Spike length

The spike length of the wheat plants varied significantly among the treatments (Fig 9I), with the test samples (7.85 ± 0.18 cm) showing intermediate values between those of the positive (8.61 ± 0.15 cm) and negative controls (6.32 ± 0.21 cm). Statistical analysis revealed that the test samples produced significantly longer spikes than did the negative control samples (p < 0.05), representing a 24.2% increase, while remaining significantly shorter than the positive control samples (8.8% reduction, p < 0.05). Compared with negative control, positive control produced 36.2% longer spikes.

**Figure 9:**
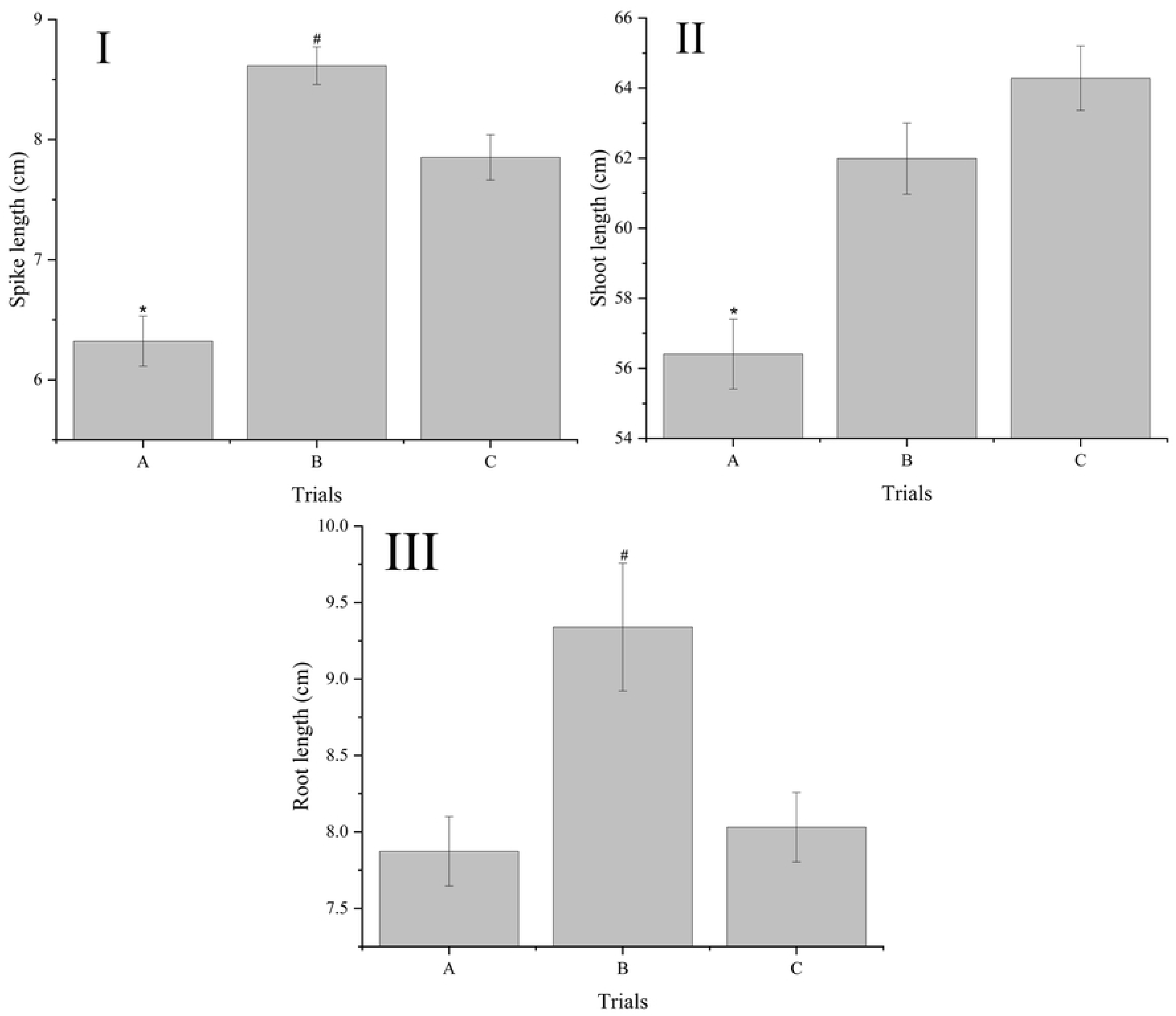
Growth parameters: (I) Spike length, (II) Shoot length, and (III) Root length. (A): Negative control, (B): DAP-supplemented positive control, (C): ST3M-39a inoculated test samples. *Significantly lower than that in trial C (α = 0.05). ^#^Significantly higher than that in trial C (α = 0.05). The data represent means ± SEs (n = 15).

##### 3.6.2.2 Shoot length

The shoot length measurements revealed significant differences among the treatments (Fig 9II). The test samples presented greater shoot elongation (64.28 ± 0.92 cm), which significantly exceeded that of the negative control (56.40 ± 0.99 cm) by 14.0% (p < 0.05). The test samples were not significantly different from the positive control (61.98 ± 1.01 cm) (3.7% decrease, p < 0.05), indicating comparable shoot growth between these treatments. Compared with the negative control, the positive control resulted in 9.9% greater shoot length.

##### 3.6.2.3 Root length

The root length measurements significantly varied among the treatments (Fig 9III), with values ranging from 7.87 to 9.34 cm. The positive control resulted in significantly greater root development (9.34 ± 0.42 cm), exceeding that of the test samples (8.03 ± 0.22 cm) by 16.3% and that of the negative control (7.87 ± 0.22 cm) by 18.7% (p < 0.05). The test samples were not significantly different from the negative control samples (2.0% increase, p > 0.05), indicating comparable root growth between these treatments.

## 4 Discussion

*Burkholderia gladioli* ST3M-39a—isolated from the maize rhizosphere and taxonomically validated through integrated morphological, biochemical, and molecular analyses—exhibits robust plant growth-promoting traits with notable phosphate solubilization capacity, positioning it as a promising bioinoculant for sustainable agriculture. Phylogenetic clustering with reference *B. gladioli* strains and 16S rRNA sequence similarity (BLAST), coupled with phenotypic congruence (urease/oxidase production, growth at pH 8/7% NaCl, utilization of sucrose/citrate/L-arginine/L-lysine/L-threonine/L-tryptophan; Table 1) absent in related species (*B. plantarii*, *B. glumae*, *B. perseverans*) (40,48,49), confirms its classification. Divergent catabolism of lactose/mannitol/pyruvate/L-cystein/glycine versus all compared species suggested strain-specific adaptations. This places ST3M-39a within well-documented *Burkholderia* genus—a dominant phosphate-solubilizing taxon alongside *Pseudomonas* and *Bacillus* (50–52) —previously identified in maize rhizospheres (25,53–55). Beyond phosphate solubilization, PSB enhances plant growth via phytohormones: auxins (root formation/cell elongation), cytokinins (cell division/senescence delay), and gibberellins (germination/stem elongation/flowering regulation) (56), consistent with reports of *B*. *gladioli* strains exhibiting ammonia production, siderophore activity, and significant plant growth promotion (25,57,58).

**Table 1:** Phenotypic features of selected isolates and comparisons with closely related species based on BLAST result.

| Test | ST3M-39a | <u>*B. gladioli*</u><br>(40) | <u>*B. perseverans*</u><br>(40)(48) | <u>*B. glumae*</u><br>(40)(49) | <u>*B. plantarii*</u><br>(40)(49) |
| --- | --- | --- | --- | --- | --- |
| **Colony morphology** |  |  |  |  |  |
| Colony form | Circular |  | Round |  |  |
| Elevation | Raised |  | Low convex |  |  |
| Margin | Entire |  |  |  |  |
| Texture | Smooth |  | Smooth |  |  |
| Opacity | Opaque |  |  |  |  |
| Surface appearance | Glistening |  | Translucent |  |  |
| Chromogenesis | Creamy<br>White |  |  |  |  |
| Diameter(mm) | 1.5 mm |  | 1mm |  |  |
| Pigmentation | - |  |  |  |  |
| **Microscopy** | gram-negative<br>rods | gram-negative,<br>non-spore<br>forming rods | gram-negative,<br>non-spore<br>forming rods |  |  |
| **VP Test** | - |  |  |  |  |
| **Indole Production** | - |  |  |  |  |
| **Motility** | - |  |  |  |  |
| **Hydrogen Sulphide Production** | - |  |  |  |  |
| **Production of:** |  |  |  |  |  |
| C4 lipase | + | + | + | v- | + |
| Amylase | - | v | v+ | v | v |
| Gelatinase | + | v+ | + | + | + |
| Protease | + | v+ | v+ | v- | + |
| Cellulase | + |  |  |  |  |
| Urease | + |  | - |  |  |
| **Growth in LB media** | + |  |  |  |  |
| **Growth in MacConkey Agar** | Pink colonies with no ppt around | - | + | - | - |
| **Oxidative in OF medium** | + |  |  |  |  |
| **Growth at** |  |  |  |  |  |
| 37°C | + |  | + |  |  |
| 40°C | + | + | + | + | + |
| 45°C | - |  |  |  |  |
| pH 8 | + | + | - | + | v- |
| 7% NaCl | + | + | - | - | - |
| **Catalase** | + |  | + |  |  |
| **Oxidase** | - | v | + |  |  |
| **Utilization of:** |  |  |  |  |  |
| Glucose | + | + | + |  |  |
| Sucrose | + | + |  | - | ± |
| Fructose | + | + |  | + | + |
| Maltose | - | - | - | - | - |
| Xylose | + | + |  | + | + |
| Galactose | + |  |  |  |  |
| Lactose | + | - | - | - | - |
| Mannitol | - | + | + |  |  |
| Melezitose | + |  |  |  |  |
| Sorbitol | + | + |  | + | + |
| Ethanol | + | + |  |  |  |
| Propanol | + | + |  |  |  |
| N-hexadecane | + |  |  |  |  |
| Sodium-K-Tartrate | + | + |  |  |  |
| Citrate | + | + | - | + | + |
| Pyruvate | - | + |  |  |  |
| L-Arginine | + | + | - | + | v |
| L-Cysteine | - | + |  | + | + |
| Glycine | + | - |  | - | - |
| L-Histidine | + | + |  | + | + |
| L-Lysine | + | + |  | - | - |
| L-Ornithine | - | + |  | - | - |
| L-Phenylalanine | + | + |  | + | + |
| L-Threonine | + | + |  | + | - |
| L-Tryptophan | + | + |  | + | - |
| L-Glutamine | + | + |  | + | + |
| **Resistant to antibiotics** | AMP, P,<br>CTX, VA,<br>RIF, CXM,<br>CB |  |  |  |  |
| **Sensitive to antibiotics** | CIP, GEN,<br>MO, LE,<br>AZM, E |  |  |  |  |

v – variable, AMP – ampicillin (10 mcg); P – penicillin (1 unit); CTX – cefotaxime (30 mcg); VA – vancomycin (5 mcg); RIF – rifampicin (5 mcg); CXM – cefuroxime (30 mcg); CB – carbenicillin (100 mcg); CIP – ciprofloxacin (5 mcg); GEN – gentamycin (10 mcg); MO – moxifloxacin (5 mcg); LE – levofloxacin (5 mcg); AZM – azithromycin (15 mcg); E – erythromycin (15 mcg).

Phosphate solubilization by ST3M-39a was confirmed through halo zone formation on PVK and NBRIP agar, with quantitative efficiency (177.96 µg/mL in liquid NBRIP) exceeding *Serratia nematodiphila* RJ10 (86 µg/mL) (59) but lower than *Burkholderia* spp. (485 - 580 mg/L at 72 h) (60) and *B. gladioli* MEL01 (350 mg/L at 28 h) (31). Its solubilization index (PSI = 1.27 PVK; PSI = 1.41 NBRIP) aligns with established media ranges (61–64), while near-equivalence to *Pseudomonas* sp. (183.2 mg/L, 72 h) (65) under shorter incubation suggests superior kinetics. These interspecies variations likely reflect divergent metabolic adaptations (52). Concomitant acidification (pH 6.8 to 5.0) confirmed organic acid-mediated solubilization, primarily via malic, gluconic, oxalic, and citric acids (22,66,67), that chelates Fe³⁺, Al³⁺, Ca²⁺ ions. Resulting stable metal-organic complexes prevents phosphate re-precipitation, enhancing phosphorus bioavailability (12). Crucially, while abiotic stresses (pH fluctuations, salinity, osmotic stress) generally constrain microbial solubilization, ST3M-39a maintained robust growth and phosphate dissolution efficiency across all tested conditions.

ST3M-39a exhibited exceptional climate resilience, maintaining growth across extreme abiotic gradients (pH 5-11, 8.5% NaCl, 0.943 a_w_)—conditions inhibiting most microorganisms (only 16.7% of arid-environment isolates survive at a_w_ 0.897 (68,69)). Previous research indicates that certain plant growth-promoting rhizobacteria (PGPR) strains exhibit robust growth across broad abiotic stress ranges, including pH extremes (4.0–12.0), salinity (2.5–7.5% NaCl), and osmotic stress induced by 20% PEG 6000 (70). ST3M-39a’s phosphate solubilization, while optimal at neutral pH without stressors (177.96 µg/mL), persisted under suboptimal conditions (pH 4.5-8.5, 7.5% NaCl, 0.950 a_w_), aligning with stress-adapted PGPR like *Priestia megaterium* (71). This pattern also aligns with previous observations of reduced but maintained phosphate solubilization under osmotic stress (59). This tolerance likely involves *Burkholderia*-characteristic adaptations: membrane remodeling, stress-protective molecules (72), EPS-mediated mechanisms. Secreted EPS mitigates salinity by chelating Na⁺ ions (73), while enhancing phosphate solubilization through dual pathways: (1) stabilizing dissolved phosphorus via carboxyl/phenolic hydroxyl groups that bind phosphate through ion exchange and hydrogen bonding (74), and (2) electrostatically repelling H⁺ from organic acids at mineral interfaces to regulate dissolution (75,76). Critically, EPS sustains phosphorus nutrition at a_w_ 0.950—when most microbial activity ceases (77)—while protecting plant from stress damage (78). This integrated stress adaptation and nutrient cycling capacity exceeds strains like *B. seminalis* 869T2 (79), positioning ST3M-39a as a premier bioinoculant candidate for climate-affected regions with saline, arid, or pH-extreme soils.

Strain ST3M-39a demonstrated a multifaceted PGPR profile, integrating phosphate/zinc solubilization, exopolysaccharide (EPS) production, protease/cellulase activity, and ammonia generation to collectively enhance plant nutrition and stress resilience. Its zinc oxide solubilization capacity matches established PGPR strains (42,80,81), Countering deficiency-induced impairments in carbohydrate metabolism, auxin synthesis, and antioxidant defense that cause chlorosis, stunted growth, and yield loss (82). EPS production, the hallmark of *Burkholderia* species (47,83,84), enhances salt tolerance through improved osmoregulation and biomass accumulation (85). Concurrently, protease and cellulase enzymes mediate dual functions: biocontrol via fungal cell wall degradation (proteolytic cleavage) and plant cell wall breakdown (hydrolysis of 1,4-β-D-glycosidic linkages), alongside nutrient mobilization (86,87); Protease further reinforce stress tolerance through protein repair, metabolic regulation, and defense hormone modulation (88,89). Finally, microbial-derived ammonia provides bioavailable nitrogen, driving root/shoot elongation and biomass increase (90).

Pot experiment demonstrated that *B. gladioli* ST3M-39a inoculation significantly enhanced wheat chlorophyll content, spike length, shoot elongation, and yield versus non-inoculated controls (Figure 8 and 9) Aligning with established evidence that PSB improve plant growth, tissue phosphorus, and yield while reducing post-harvest soil organic matter and pH (91–95). Although PSB-inoculated growth significantly exceeded negative control (p < 0.05), it remained below DAP-fertilized plants, reflecting increased—yet comparatively limited—bioavailable soil phosphorus relative to soluble DAP (96). Crucially, ST3M-39a achieved 82-90% of DAP’s fertilizer efficiency while significantly enhancing phosphorus uptake, demonstrating viability as a multifunctional biofertilizer whose integrated traits (phosphate/zinc solubilization, EPS production, protease/cellulase activity, ammonia production) collectively boost nutrient bioavailability and growth (97). Field data confirmed yield increased with reduced chemical inputs. While current microbial solubilization kinetics constrain parity with chemical phosphorus sources, this gap is addressable through targeted optimization of organic acid pathways and hybrid systems combining ST3M-39a with reduced DAP (98); strategic strain engineering and field validation could position it as a complete alternative for climate-vulnerable agriculture within a decade (99), delivering agronomic equivalence while enhancing long-term soil health via improved nutrient cycling (100).

## 5 Conclusion

*Burkholderia gladioli* ST3M-39a emerges as a transformative bioinoculant, demonstrating exceptional resilience to abiotic stresses, while maintaining robust phosphate solubilization (177.96 ± 5.26 µg/mL). Its multifunctional plant growth-promoting traits (zinc solubilization, EPS production, enzymatic activity, and ammonia synthesis) drove wheat yield and biomass to 85-92% of fertilizer-equivalent yields under controlled conditions. This convergence of climate adaptability and nutrient mobilization efficiency positions ST3M-39a as a scalable solution for revitalizing phosphorus-deficient marginal soils.

To translate this potential into tangible agricultural impact, farmers should pilot field applications via seed coating or soil drenching, particularly in saline/acidic/drought-affected zones, blending inoculants with 50% reduced synthetic fertilizers to lower costs. Policymakers must prioritize funding local production hubs and integrating bioinoculants into subsidy programs, contingent on rigorous 2-year ecological safety assessments. Researchers ought to accelerate multi-location trials to validate performance across heterogeneous field conditions, develop self-stable formulations, and monitor long-term soil microbiome impacts. Collaborative efforts among agronomists, microbiologists, and policymakers are now critical to deploy ST3M-39a as a renewable alternative, advancing toward the restoration of degraded global croplands.

## 6 Acknowledgement

We gratefully acknowledge Shubham Biotech Nepal Private Limited for providing essential research space and facilities. Special thanks are extended to the National Biotechnology Research Center, National Agricultural Research Council for their support and provision of greenhouse facilities. We also express sincere appreciation to Mr. Dipak Raj Pandey and Mr. Anish Basnet for their invaluable assistance throughout this study.

## 7 Data availability

The datasets generated in this study are available in the Mendeley Data repository (DOI: 10.17632/h7g7cp3jxk.1) (101). The 16S rRNA nucleotide sequence is deposited in GenBank under accession number OR143748.1.

## Reference

1. Kaur T, Manhas RK. Evaluation of ACC deaminase and indole acetic acid production by Streptomyces hydrogenans DH16 and its effect on plant growth promotion. Biocatal Agric Biotechnol [Internet]. 2022 Jul;42(March):102321. Available from: 10.1016/j.bcab.2022.102321

2. Gupta A, Bano A, Rai S, Kumar M, Ali J, Sharma S, Pathak N. ACC deaminase producing plant growth promoting rhizobacteria enhance salinity stress tolerance in Pisum sativum. 3 Biotech [Internet]. 2021 Dec 29;11(12):514. Available from: https://link.springer.com/10.1007/s13205-021-03047-5

3. Orozco-Mosqueda MDC, Glick BR, Santoyo G. ACC deaminase in plant growth-promoting bacteria (PGPB): An efficient mechanism to counter salt stress in crops. Microbiol Res [Internet]. 2020 May;235(January):126439. Available from: http://www.ncbi.nlm.nih.gov/pubmed/32097862

4. Orozco-Mosqueda MDC, Glick BR, Santoyo G. IAA and ACC deaminase producing-bacteria isolated from the rhizosphere of pineapple plants grown under different abiotic and biotic stresses. Microbiol Res [Internet]. 2020 May;235(6):126439. Available from: http://www.ncbi.nlm.nih.gov/pubmed/32097862

5. Siedliska A, Baranowski P, Pastuszka-Woźniak J, Zubik M, Krzyszczak J. Identification of plant leaf phosphorus content at different growth stages based on hyperspectral reflectance. BMC Plant Biol. 2021;21(1):1–17. Available from: 10.1186/s12870-020-02807-4

6. Chouyia FE, Ventorino V, Pepe O. Diversity, mechanisms and beneficial features of phosphate-solubilizing Streptomyces in sustainable agriculture: A review. Front Plant Sci. 2022;13(December):1–16. Available from: 10.3389/fpls.2022.1035358

7. Elhaissoufi W, Ghoulam C, Barakat A, Zeroual Y, Bargaz A. Phosphate bacterial solubilization: A key rhizosphere driving force enabling higher P use efficiency and crop productivity. J Adv Res [Internet]. 2022;38:13–28. Available from: 10.1016/j.jare.2021.08.014

8. Suleimanova AD, Beinhauer A, Valeeva LR, Chastukhina IB, Balaban NP, Shakirov E V., Greiner R, Sharipova MR. Novel glucose-1-phosphatase with high phytase activity and unusual metal ion activation from soil bacterium *Pantoea* sp. Strain 3.5.1. Elliot MA, editor. Appl Environ Microbiol [Internet]. 2015 Oct;81(19):6790–9. Available from: https://journals.asm.org/doi/10.1128/AEM.01384-15

9. Pan L, Cai B. Phosphate-solubilizing bacteria: advances in their physiology, molecular mechanisms and microbial community effects. Microorganisms [Internet]. 2023 Dec 1;11(12):2904. Available from: https://www.mdpi.com/2076-2607/11/12/2904

10. Santos HL, Silva GF da, Carnietto MRA, Oliveira LC, Nogueira CH de C, Silva M de A. *Bacillus velezensis* Associated with organomineral fertilizer and reduced phosphate doses improves soil microbial—chemical properties and biomass of sugarcane. Agronomy. 2022;12(11). Available from: 10.3390/agronomy12112701

11. Singh AK, Singh JB, Singh R, Kantwa SR, Jha PK, Ahamad S, Singh A, Ghosh A, Prasad M, Singh S, Singh S, Prasad PVV. Understanding soil carbon and phosphorus dynamics under grass-legume intercropping in a semi-arid region. Agronomy. 2023;13(7). Available from: 10.3390/agronomy13071692

12. Ahash S, Manikandan K, Sivasankari Devi T, Elamathi S, Maragatham S, Subrahmaniyan K. Phosphate-solubilizing microorganisms for sustainable phosphorus management in rice. Rhizosphere [Internet]. 2025;34:101096. Available from: https://www.sciencedirect.com/science/article/pii/S2452219825000813

13. Mei C, Chretien RL, Amaradasa BS, He Y, Turner A, Lowman S. Characterization of phosphate solubilizing bacterial endophytes and plant growth promotion in vitro and in greenhouse. Microorganisms [Internet]. 2021 Sep 11;9(9):1935. Available from: https://www.mdpi.com/2076-2607/9/9/1935

14. Wendimu A, Yoseph T, Ayalew T. Ditching phosphatic fertilizers for phosphate-solubilizing biofertilizers: A step towards sustainable agriculture and environmental health. Sustainability [Internet]. 2023 Jan 16;15(2):1713. Available from: https://www.mdpi.com/2071-1050/15/2/1713

15. Illakwahhi DT, Vegi MR, Srivastava BBL. Phosphorus’ future insecurity, the horror of depletion, and sustainability measures. Int J Environ Sci Technol [Internet]. 2024 Oct 8;21(14):9265–80. Available from: 10.1007/s13762-024-05664-y

16. Gross A, Lin Y, Weber PK, Pett-Ridge J, Silver WL. The role of soil redox conditions in microbial phosphorus cycling in humid tropical forests. Ecology [Internet]. 2020 Feb 17;101(2):1–10. Available from: 10.1038/s41396-020-0632-4

17. Mohamed AE, Nessim MG, Abou-el-seoud II, Darwish KM, Shamseldin A. Isolation and selection of highly effective phosphate solubilizing bacterial strains to promote wheat growth in Egyptian calcareous soils. Bull Natl Res Cent [Internet]. 2019 Dec 27;43(1):203. Available from: https://bnrc.springeropen.com/articles/10.1186/s42269-019-0212-9

18. Li Y, Liu X, Hao T, Chen S. Colonization and maize growth promotion induced by phosphate solubilizing bacterial isolates. Int J Mol Sci. 2017;18(7). Available from: 10.3390/ijms18071253

19. Chawngthu L, Hnamte R, Lalfakzuala R. Isolation and characterization of rhizospheric phosphate solubilizing bacteria from wetland paddy field of Mizoram, India. Geomicrobiol J [Internet]. 2020 Apr 20;37(4):366–75. Available from: 10.1080/01490451.2019.1709108

20. Bhakat K, Chakraborty A, Islam E. Characterization of zinc solubilization potential of arsenic tolerant *Burkholderia* spp. isolated from rice rhizospheric soil. World J Microbiol Biotechnol [Internet]. 2021;37(3):1–13. Available from: 10.1007/s11274-021-03003-8

21. Wang Z, Zhang H, Liu L, Li S, Xie J, Xue X, Jiang Y. Screening of phosphate-solubilizing bacteria and their abilities of phosphorus solubilization and wheat growth promotion. BMC Microbiol [Internet]. 2022 Dec 9;22(1):296. Available from: 10.1186/s12866-022-02715-7

22. Rawat P, Das S, Shankhdhar D, Shankhdhar SC. Phosphate-solubilizing microorganisms: mechanism and their role in phosphate solubilization and uptake. J Soil Sci Plant Nutr [Internet]. 2021 Mar 30;21(1):49–68. Available from: https://link.springer.com/10.1007/s42729-020-00342-7

23. Saeed Q, Xiukang W, Haider FU, Kučerik J, Mumtaz MZ, Holatko J, Naseem M, Kintl A, Ejaz M, Naveed M, Brtnicky M, Mustafa A. Rhizosphere bacteria in plant growth promotion, biocontrol, and bioremediation of contaminated sites: A comprehensive review of effects and mechanisms. Int J Mol Sci [Internet]. 2021 Sep 29;22(19):10529. Available from: https://www.mdpi.com/1422-0067/22/19/10529

24. Gómez-Godínez LJ, Aguirre-Noyola JL, Martínez-Romero E, Arteaga-Garibay RI, Ireta-Moreno J, Ruvalcaba-Gómez JM. A look at plant-growth-promoting bacteria. Plants [Internet]. 2023 Apr 17;12(8):1668. Available from: http://ieeexplore.ieee.org/document/4625413/

25. Guo D-J, Yang G-R, Singh P, Wang J-J, Lan X-M, Singh RK, Guo J, Dong Y, Li D, Yang B. Comprehensive analysis of the physiological and molecular responses of phosphate-solubilizing bacterium *Burkholderia gladioli* DJB4–8 in promoting maize growth. Front Plant Sci [Internet]. 2025 Jun 13;16(June):1–20. Available from: https://www.frontiersin.org/articles/10.3389/fpls.2025.1611674/full

26. Pal G, Saxena S, Kumar K, Verma A, Sahu PK, Pandey A, White JF, Verma SK. Endophytic *Burkholderia*: Multifunctional roles in plant growth promotion and stress tolerance. Microbiol Res [Internet]. 2022 Dec;265(July):127201. Available from: 10.1016/j.micres.2022.127201

27. Bach E, Passaglia LMP, Jiao J, Gross H. *Burkholderia* in the genomic era: from taxonomy to the discovery of new antimicrobial secondary metabolites. Crit Rev Microbiol [Internet]. 2022 Mar 4;48(2):121–60. Available from: 10.1080/1040841X.2021.1946009

28. Shao J, Miao Y, Liu K, Ren Y, Xu Z, Zhang N, Feng H, Shen Q, Zhang R, Xun W. Rhizosphere microbiome assembly involves seed-borne bacteria in compensatory phosphate solubilization. Soil Biol Biochem [Internet]. 2021 Aug;159(May):108273. Available from: 10.1016/j.soilbio.2021.108273

29. Mannaa M, Park I, Seo Y-S. Genomic features and insights into the taxonomy, virulence, and benevolence of plant-associated *Burkholderia* Species. Int J Mol Sci [Internet]. 2018 Dec 29;20(1):121. Available from: https://www.mdpi.com/1422-0067/20/1/121

30. Khiangte L, Lalfakzuala R. Effects of heavy metals on phosphatase enzyme activity and indole-3-acetic acid (IAA) production of phosphate solubilizing bacteria. Geomicrobiol J [Internet]. 2021 Jun 1;38(6):494–503. Available from: 10.1080/01490451.2021.1894271

31. Sun X, Shao C, Chen L, Jin X, Ni H. Plant growth-promoting effect of the chitosanolytic phosphate-solubilizing bacterium *Burkholderia gladioli* MEL01 after fermentation with chitosan and fertilization with rock phosphate. J Plant Growth Regul [Internet]. 2021 Aug 23;40(4):1674–86. Available from: 10.1007/s00344-020-10223-z

32. Koh CM. Storage of bacteria and yeast [Internet]. 1st ed. Vol. 533, Methods in Enzymology. Elsevier Inc.; 2013. 15–21 p. Available from: 10.1016/B978-0-12-420067-8.00002-7

33. Pikovskaya RI, Pikovskaya RI. Mobilization of phosphorus in soil in connection with the vital activity of some microbial species. In 1948. Available from: https://www.cabidigitallibrary.org/doi/full/10.5555/19491901720

34. Nautiyal CS. An efficient microbiological growth medium for screening phosphate solubilizing microorganisms. FEMS Microbiol Lett. 1999;170(1):265–70. Available from: 10.1016/S0378-1097(98)00555-2

35. Murphy J, Riley JP. A modified single solution method for the determination of phosphate in natural waters. Anal Chim Acta [Internet]. 1962;27(C):31–6. Available from: https://linkinghub.elsevier.com/retrieve/pii/S0003267000884445

36. Sambrook J, Russell DW. Molecular cloning : a laboratory manual [Internet]. 3rd ed. COLD SPRING HARBOR LABORATORY PRESS. Cold Spring Harbor, N.Y. SE -: Cold Spring Harbor Laboratory Press; 2001. Available from: https://www.cshlpress.com/pdf/sample/2013/MC4/MC4FM.pdf

37. Tamura K, Stecher G, Kumar S. MEGA11: Molecular evolutionary genetics analysis version 11. Battistuzzi FU, editor. Mol Biol Evol [Internet]. 2021 Jun 25;38(7):3022–7. Available from: https://academic.oup.com/mbe/article/38/7/3022/6248099

38. Tamura K, Nei M. Estimation of the number of nucleotide substitutions in the control region of mitochondrial DNA in humans and chimpanzees. Mol Biol Evol [Internet]. 1993 May;10(3):512–26. Available from: https://academic.oup.com/mbe/article/10/3/512/1016366/Estimation-of-the-number-of-nucleotide

39. Felsenstein J. Confidence limits on phylogenies: An approach using the bootstrap. evolution (N Y) [Internet]. 1985 Jul;39(4):783. Available from: https://www.jstor.org/stable/2408678?origin=crossref

40. Brenner DJ, Krieg NR, Garrity GM, Staley JTTA-TT-. Bergey’s manual of systematic bacteriology [Internet]. 2nd ed NV. Brenner DJ, Krieg NR, Staley JT, Garrity GM, editors. Boston, MA: Springer US; 2005. Available from: http://link.springer.com/10.1007/0-387-28021-9

41. Fysun O, Stoeckel M, Thienel KJF, Wäschle F, Palzer S, Hinrichs J. Prediction of water activity in aqueous polyol solutions. Chemie Ing Tech [Internet]. 2015 Oct 11;87(10):1327–33. Available from: https://onlinelibrary.wiley.com/doi/10.1002/cite.201400134

42. Gontia-Mishra I, Sapre S, Tiwari S. Zinc solubilizing bacteria from the rhizosphere of rice as prospective modulator of zinc biofortification in rice. Rhizosphere [Internet]. 2017 Jun;3:185–90. Available from: 10.1016/j.rhisph.2017.04.013

43. Khan N, Bano A. Exopolysaccharide producing rhizobacteria and their impact on growth and drought tolerance of wheat grown under rainfed conditions. Singh AK, editor. PLoS One [Internet]. 2019 Sep 12;14(9):e0222302. Available from: 10.1371/journal.pone.0222302

44. Latif M, Bukhari SAH, Alrajhi AA, Alotaibi FS, Ahmad M, Shahzad AN, Dewindar AZ, Mattar MA. Inducing drought tolerance in wheat through exopolysaccharide-producing rhizobacteria. Agronomy [Internet]. 2022 May 9;12(5):1140. Available from: https://www.mdpi.com/2073-4395/12/5/1140

45. Demissie MS, Legesse NH, Tesema AA. Isolation and characterization of cellulase producing bacteria from forest, cow dung, Dashen brewery and agro-industrial waste. PLoS One [Internet]. 2024;19(4 April):1–12. Available from: 10.1371/journal.pone.0301607

46. Pandey GR, Shrestha A, Karki TB, Neupane S, Ojha S, Koirala P, Timilsina PM. Screening and identification of thermotolerant and osmotolerant bacillus amyloliquefaciens bkhe isolated from kinema of eastern nepal for alkaline protease production. Li Z, editor. Int J Microbiol [Internet]. 2022 Dec 6;2022:1–11. Available from: https://www.hindawi.com/journals/ijmicro/2022/6831092/

47. Chen E, Yang C, Tao W, Li S. Polysaccharides produced by plant growth-promoting rhizobacteria strain *Burkholderia* sp. BK01 enhance salt stress tolerance to *Arabidopsis thaliana*. Polymers (Basel) [Internet]. 2024 Jan 3;16(1):145. Available from: https://www.mdpi.com/2073-4360/16/1/145

48. Andrade JP, de Souza HG, Ferreira LC, Cnockaert M, De Canck E, Wieme AD, Peeters C, Gross E, De Souza JT, Marbach PAS, Góes-Neto A, Vandamme P. *Burkholderia perseverans* sp. nov., a bacterium isolated from the Restinga ecosystem, is a producer of volatile and diffusible compounds that inhibit plant pathogens. Braz J Microbiol [Internet]. 2021 Dec 21;52(4):2145–52. Available from: 10.1007/s42770-021-00560-w

49. Viallard V, Poirier I, Cournoyer B, Haurat J, Wiebkin S, Ophel-Keller K, Balandreau J. *Burkholderia graminis* sp. nov., a rhizospheric *Burkholderia* species, and reassessment of [*Pseudomonas*] *phenazinium*, [*Pseudomonas*] *pyrrocinia* and [*Pseudomonas*] *glathei* as *Burkholderia*. Int J Syst Bacteriol [Internet]. 1998 Apr 1;48(2):549–63. Available from: https://www.microbiologyresearch.org/content/journal/ijsem/10.1099/00207713-48-2-549

50. Wang C, Pan G, Lu X, Qi W. Phosphorus solubilizing microorganisms: potential promoters of agricultural and environmental engineering. Front Bioeng Biotechnol [Internet]. 2023 May 12;11(May):1–5. Available from: https://www.frontiersin.org/articles/10.3389/fbioe.2023.1181078/full

51. Song C, Wang W, Gan Y, Wang L, Chang X, Wang Y, Yang W. Growth promotion ability of phosphate-solubilizing bacteria from the soybean rhizosphere under maize–soybean intercropping systems. J Sci Food Agric [Internet]. 2022 Mar 15;102(4):1430–42. Available from: 10.1002/jsfa.11477

52. Chen J, Zhao G, Wei Y, Dong Y, Hou L, Jiao R. Isolation and screening of multifunctional phosphate solubilizing bacteria and its growth-promoting effect on Chinese fir seedlings. Sci Rep [Internet]. 2021 Apr 27;11(1):9081. Available from: 10.1038/s41598-021-88635-4

53. Mohd Din ARJ, Othman NZ. Genome sequence data of *Burkholderia* sp. IMCC1007 isolated from maize rhizosphere: A potential strain in fusaric acid mycotoxin biodegradation. Data Br [Internet]. 2023 Jun;48:109204. Available from: 10.1016/j.dib.2023.109204

54. Medina-de la Rosa G, López-Reyes L, Carcaño-Montiel MG, López-Olguín JF, Hernández-Espinosa MÁ, Rivera-Tapia JA. Rhizosphere bacteria of maize with chitinolytic activity and its potential in the control of phytopathogenic fungi. Arch Phytopathol Plant Prot [Internet]. 2016 Jul 20;49(11–12):310–21. Available from: https://www.tandfonline.com/doi/full/10.1080/03235408.2016.1201345

55. Silva K da, Perin L, Gomes M de L, Baraúna AC, Pereira GMD, Mosqueira CA, Costa IB da, O’hara G, Zilli J É. Diversity and capacity to promote maize growth of bacteria isolated from the Amazon region. Acta Amaz [Internet]. 2016 Jun;46(2):111–8. Available from: http://www.scielo.br/scielo.php?script=sci_arttext&pid=S0044-59672016000200111&lng=en&tlng=en

56. Puri A, Padda KP, Chanway CP. In vitro and in vivo analyses of plant-growth-promoting potential of bacteria naturally associated with spruce trees growing on nutrient-poor soils. Appl Soil Ecol [Internet]. 2020 May;149(December 2019):103538. Available from: 10.1016/j.apsoil.2020.103538

57. Li Z, Li J, Liu G, Li Y, Wu X, Liang J, Wang Z, Chen Q, Peng F. Isolation, characterization and growth-promoting properties of phosphate-solubilizing bacteria (PSBs) derived from peach tree rhizosphere. Microorganisms [Internet]. 2025 Mar 23;13(4):718. Available from: https://www.mdpi.com/2076-2607/13/4/718

58. Fitriatin BN, Mulyani O, Herdiyantoro D, Alahmadi TA, Pellegrini M. Metabolic characterization of phosphate solubilizing microorganisms and their role in improving soil phosphate solubility, yield of upland rice (*Oryza sativa L*.), and phosphorus fertilizers efficiency. Front Sustain Food Syst [Internet]. 2022 Nov 10;6. Available from: https://www.frontiersin.org/articles/10.3389/fsufs.2022.1032708/full

59. Saikia J, Sarma RK, Dhandia R, Yadav A, Bharali R, Gupta VK, Saikia R. Alleviation of drought stress in pulse crops with ACC deaminase producing rhizobacteria isolated from acidic soil of Northeast India. Sci Rep [Internet]. 2018 Feb 23;8(1):3560. Available from: https://www.nature.com/articles/s41598-018-21921-w

60. Ghosh R, Barman S, Mukherjee R, Mandal NC. Role of phosphate solubilizing Burkholderia spp. for successful colonization and growth promotion of *Lycopodium cernuum L*. (*Lycopodiaceae*) in lateritic belt of Birbhum district of West Bengal, India. Microbiol Res [Internet]. 2016 Feb;183:80–91. Available from: 10.1016/j.micres.2015.11.011

61. Gupta R, Kumari A, Sharma S, Alzahrani OM, Noureldeen A, Darwish H. Identification, characterization and optimization of phosphate solubilizing rhizobacteria (PSRB) from rice rhizosphere. Saudi J Biol Sci [Internet]. 2022 Jan;29(1):35–42. Available from: 10.1016/j.sjbs.2021.09.075

62. Mengesha AS, Legesse NH. Isolation and characterization of phosphate solubilizing bacteria from the rhizosphere of lentil *(Lens culinaris M*.) collected from Hagere Mariam district, Central Ethiopia. Kee Zuan AT, editor. PLoS One [Internet]. 2024 Nov 15;19(11):e0308915. Available from: 10.1371/journal.pone.0308915

63. Rfaki A, Zennouhi O, Aliyat FZ, Nassiri L, Ibijbijen J. Isolation, selection and characterization of root-associated rock phosphate solubilizing bacteria in Moroccan wheat ( *Triticum aestivum L*.). Geomicrobiol J [Internet]. 2020 Mar 15;37(3):230–41. Available from: 10.1080/01490451.2019.1694106

64. Manzoor M, Abbasi MK, Sultan T. Isolation of phosphate solubilizing bacteria from maize rhizosphere and their potential for rock phosphate solubilization–mineralization and plant growth promotion. Geomicrobiol J [Internet]. 2017 Jan 2;34(1):81–95. Available from: https://www.tandfonline.com/doi/full/10.1080/01490451.2016.1146373

65. Cheng Y, Guo M, Fan R, Chen H, Liu X, Zhu J, Wei C, Huang S, Zhang T. Screening and characterization of highly efficient phosphorus solubilizing bacteria in ecologically fragile areas. J Phys Conf Ser [Internet]. 2025 May 1;3015(1):012006. Available from: https://iopscience.iop.org/article/10.1088/1742-6596/3015/1/012006

66. Su M, Han F, Wu Y, Yan Z, Lv Z, Tian D, Wang S, Hu S, Shen Z, Li Z. Effects of phosphate-solubilizing bacteria on phosphorous release and sorption on montmorillonite. Appl Clay Sci [Internet]. 2019 Nov;181(July):105227. Available from: https://linkinghub.elsevier.com/retrieve/pii/S0169131719302856

67. Sarmah R, Sarma AK. Phosphate solubilizing microorganisms: A review. Commun Soil Sci Plant Anal [Internet]. 2023 May 31;54(10):1306–15. Available from: 10.1080/00103624.2022.2142238

68. Stevenson A, Cray JA, Williams JP, Santos R, Sahay R, Neuenkirchen N, McClure CD, Grant IR, Houghton JD, Quinn JP, Timson DJ, Patil SV, Singhal RS, Antón J, Dijksterhuis J, Hocking AD, Lievens B, Rangel DEN, Voytek MA, GundeCimerman N, Oren A, Timmis KN, McGenity TJ, Hallsworth JE. Is there a common water-activity limit for the three domains of life? ISME J [Internet]. 2015 Jun 1;9(6):1333–51. Available from: https://academic.oup.com/ismej/article/9/6/1333-1351/7558135

69. Shreshtha K, Prakash A, Pandey PK, Pal AK, Singh J, Tripathi P, Mitra D, Jaiswal DK, Santos-Villalobos, S. de los, Tripathi V. Isolation and characterization of plant growth promoting rhizobacteria from cacti root under drought condition. Curr Res Microb Sci [Internet]. 2025;8(November 2024):100319. Available from: 10.1016/j.crmicr.2024.100319

70. Thakur R, Rahi P, Gulati A, Gulati A. Tea seedlings growth promotion by widely distributed and stress-tolerant PGPR from the acidic soils of the Kangra valley. BMC Microbiol [Internet]. 2025 Feb 28;25(1):102. Available from: 10.1186/s12866-025-03811-0

71. Thakur R, Dhar H, Swarnkar MK, Soni R, Sharma KC, Singh A, Gulati A, Sud RK, Gulati A. Understanding the molecular mechanism of PGPR strain *Priestia megaterium* from tea rhizosphere for stress alleviation and crop growth enhancement. Plant Stress [Internet]. 2024 Jun;12(May):100494. Available from: 10.1016/j.stress.2024.100494

72. Duangurai T, Indrawattana N, Pumirat P. *Burkholderia pseudomallei* adaptation for survival in stressful conditions. Biomed Res Int [Internet]. 2018 May 27;2018:3039106. Available from: https://www.hindawi.com/journals/bmri/2018/3039106/

73. Shultana R, Kee Zuan AT, Yusop MR, Saud HM. Characterization of salt-tolerant plant growth-promoting rhizobacteria and the effect on growth and yield of saline-affected rice. Lopes AR, editor. PLoS One [Internet]. 2020 Sep 4;15(9):e0238537. Available from: 10.1371/journal.pone.0238537

74. Mattey M. The production of organic acids. Crit Rev Biotechnol [Internet]. 1992 Jan 27;12(1–2):87–132. Available from: http://www.tandfonline.com/doi/full/10.3109/07388559209069189

75. Yang W, Xu L, Wang Z, Li K, Hu R, Su J, Zhang L. Synchronous removal of ammonia nitrogen, phosphate, and calcium by heterotrophic nitrifying strain *Pseudomonas* sp. Y1 based on microbial induced calcium precipitation. Bioresour Technol [Internet]. 2022 Nov;363:127996. Available from: https://www.sciencedirect.com/science/article/pii/S0960852422013293

76. Zhou G, Jia X, Xu Y, Gao X, Zhao Z, Li L. Efficient remediation of cadmium and lead contaminated soil in coal mining areas by MICP application in hydrothermal carbon-based bacterial agents: Nucleation pathways and mineralization mechanisms. J Environ Manage [Internet]. 2024 Nov;370:122744. Available from: https://www.sciencedirect.com/science/article/pii/S0301479724027300

77. Breitkreuz C, Buscot F, Tarkka M, Reitz T. Shifts between and among populations of wheat rhizosphere *Pseudomonas*, Streptomyces and Phyllobacterium suggest consistent phosphate mobilization at different wheat growth stages under abiotic stress. Front Microbiol [Internet]. 2020 Jan 22;10(January). Available from: https://www.frontiersin.org/article/10.3389/fmicb.2019.03109/full

78. Verma H, Kumar D, Kumar V, Kumari M, Singh SK, Sharma VK, Droby S, Santoyo G, White JF, Kumar A. The potential application of endophytes in management of stress from drought and salinity in crop plants. Microorganisms [Internet]. 2021 Aug 13;9(8):1729. Available from: https://link.springer.com/10.1007/s10811-021-02570-5

79. Hwang H-H, Huang Y-T, Chien P-R, Huang F-C, Wu C-L, Chen L-Y, Hung S-HW, Pan I-C, Huang C-C. A plant endophytic bacterium *Burkholderia seminalis* strain 869T2 increases plant growth under salt stress by affecting several phytohormone response pathways. Bot Stud [Internet]. 2025 Feb 4;66(1):7. Available from: https://as-botanicalstudies.springeropen.com/articles/10.1186/s40529-025-00453-3

80. Sirohi G, Upadhyay A, Srivastava P., Srivastava S. PGPR mediated Zinc biofertilization of soil and its impact on growth and productivity of wheat. J soil Sci plant Nutr [Internet]. 2015;15(ahead):0–0. Available from: http://www.scielo.cl/scielo.php?script=sci_arttext&pid=S0718-95162015005000017&lng=en&nrm=iso&tlng=en

81. Dinesh R, Srinivasan V, Hamza S, Sarathambal C, Anke Gowda SJ, Ganeshamurthy AN, Gupta SB, Aparna NV, Subila KP, Lijina A, Divya VC. Isolation and characterization of potential Zn solubilizing bacteria from soil and its effects on soil Zn release rates, soil available Zn and plant Zn content. Geoderma [Internet]. 2018;321(February):173–86. Available from: 10.1016/j.geoderma.2018.02.013

82. Kamran S, Shahid I, Baig DN, Rizwan M, Malik KA, Mehnaz S. Contribution of zinc solubilizing bacteria in growth promotion and zinc content of wheat. Front Microbiol [Internet]. 2017 Dec 21;8(DEC). Available from: http://journal.frontiersin.org/article/10.3389/fmicb.2017.02593/full

83. dos Santos IB, Pereira AP de A, de Souza AJ, Cardoso EJBN, da Silva FG, Oliveira JTC, Verdi MCQ, Sobral JK Selection and characterization of *Burkholderia* spp. for their plant-growth promoting effects and influence on maize seed germination. Front Soil Sci [Internet]. 2022 Jan 12;1(January):1–10. Available from: https://www.frontiersin.org/articles/10.3389/fsoil.2021.805094/full

84. Barrera-Galicia GC, Peniche-Pavía HA, Peña-Cabriales JJ, Covarrubias SA, Vera-Núñez JA, Délano-Frier JP. Metabolic footprints of *Burkholderia Sensu Lato* rhizosphere bacteria active against maize *Fusarium* pathogens. Microorganisms [Internet]. 2021 Sep 29;9(10):2061. Available from: https://www.mdpi.com/2076-2607/9/10/2061

85. Sun L, Lei P, Wang Q, Ma J, Zhan Y, Jiang K, Xu Z, Xu H. The endophyte *Pantoea alhagi* NX-11 alleviates salt stress damage to rice seedlings by secreting exopolysaccharides. Front Microbiol [Internet]. 2020 Jan 22;10(January):1–13. Available from: https://www.frontiersin.org/article/10.3389/fmicb.2019.03112/full

86. Dutta P, Muthukrishnan G, Kutalingam Gopalasubramaiam S, Dharmaraj R, Karuppaiah A, Periyasamy K, Arumugam PM, Upamanya G, Boruah S, Deb I, Kumari A, Mahanta M, Heisnam P, Mishra A. Plant growth-promoting rhizobacteria (PGPR) and its mechanisms against plant diseases for sustainable agriculture and better productivity. BIOCELL [Internet]. 2022;46(8):1843–59. Available from: https://www.techscience.com/biocell/v46n8/47566

87. Wang H, Liu R, You MP, Barbetti MJ, Chen Y. Pathogen Biocontrol using plant growth-promoting bacteria (PGPR): role of bacterial diversity. Microorganisms [Internet]. 2021 Sep 18;9(9):1988. Available from: https://www.mdpi.com/2076-2607/9/9/1988

88. D’Ippólito S, Rey-Burusco MF, Feingold SE, Guevara MG. Role of proteases in the response of plants to drought. Plant Physiol Biochem [Internet]. 2021 Nov;168:1–9. Available from: https://www.sciencedirect.com/science/article/pii/S0981942821005039

89. Kazerooni EA, Maharachchikumbura SSN, Adhikari A, Al-Sadi AM, Kang S-M, Kim L-R, Lee I-J. Rhizospheric *Bacillus amyloliquefaciens* Protects *Capsicum annuum* cv. Geumsugangsan from multiple abiotic stresses via multifarious plant growth-promoting attributes. Front Plant Sci [Internet]. 2021 May 25;12(May):1–19. Available from: https://www.frontiersin.org/articles/10.3389/fpls.2021.669693/full

90. Bhattacharyya C, Banerjee S, Acharya U, Mitra A, Mallick I, Haldar A, Haldar S, Ghosh A, Ghosh A. Evaluation of plant growth promotion properties and induction of antioxidative defense mechanism by tea rhizobacteria of Darjeeling, India. Sci Rep [Internet]. 2020 Sep 23;10(1):15536. Available from: 10.1038/s41598-020-72439-z

91. Yang G, Liu C, Gu L, Chen Q, Zhang X. Studies on the phosphorus-solubilizing ability of isaria cateinannulata and its influence on the growth of *Fagopyrum tataricum* plants. Plants [Internet]. 2024 Jun 19;13(12):1694. Available from: https://www.mdpi.com/2223-7747/13/12/1694

92. Kaur S, Kalia A, Sharma S. Bioformulation of *Azotobacter* and *Streptomyces* for improved growth and yield of wheat (*Triticum aestivum L*.): A field study. J Plant Growth Regul [Internet]. 2024 Aug 16;43(8):2555–71. Available from: 10.1007/s00344-024-11282-2

93. Damo JLC, Pedro M, Sison ML. Phosphate solubilization and plant growth promotion by *Enterobacter* sp. isolate. Appl Microbiol [Internet]. 2024 Jul 31;4(3):1177–92. Available from: https://www.mdpi.com/2673-8007/4/3/80

94. Adnan M, Fahad S, Saleem MH, Ali B, Mussart M, Ullah R, Amanullah AM, Ahmad M, Shah WA, Romman M, Wahid F, Wang D, Saud S, Liu K, Harrison MT, Wu C, Danish S, Datta R, Muresan CC, Marc RA. Comparative efficacy of phosphorous supplements with phosphate solubilizing bacteria for optimizing wheat yield in calcareous soils. Sci Rep [Internet]. 2022 Jul 14;12(1):11997. Available from: 10.1038/s41598-022-16035-3

95. Khourchi S, Elhaissoufi W, Loum M, Ibnyasser A, Haddine M, Ghani R, et al. Phosphate solubilizing bacteria can significantly contribute to enhance P availability from polyphosphates and their use efficiency in wheat. Microbiol Res [Internet]. 2022 Sep;262(June):127094. Available from: 10.1016/j.micres.2022.127094

96. Torres-Cuesta D, Mora-Motta D, Chavarro-Bermeo JP, Olaya-Montes A, Vargas-Garcia C, Bonilla R, et al. Phosphate-Solubilizing bacteria with low-solubility fertilizer improve soil p availability and yield of kikuyu grass. Microorganisms [Internet]. 2023 Jul 4;11(7):1748. Available from: https://www.mdpi.com/2076-2607/11/7/1748

97. Murad S, Ahmad M, Hussain A, Ali S, Al-Ansari N, Mattar MA. Efficacy of DAP coated with bacterial strains and their metabolites for soil phosphorus availability and maize growth. Sci Rep [Internet]. 2024 May 18;14(1):11389. Available from: 10.1038/s41598-024-61817-6

98. Tang Y, Che Y-J, Bai X-Y, Wang Z-Y, Gu S-Y. Effects of application of phosphate and phosphate-solubilizing bacteria on bacterial diversity and phosphorus fractions in a Phaeozems. Heliyon [Internet]. 2023 Dec;9(12):e22937. Available from: https://linkinghub.elsevier.com/retrieve/pii/S2405844023101459

99. Benitez-Alfonso Y, Soanes BK, Zimba S, Sinanaj B, German L, Sharma V, Bohra A, Kolesnikova A, Dunn JA, Martin AC, Khashi u Rahman M, Saati-Santamaría Z, García-Fraile P, Ferreira EA, Frazão LA, Cowling WA, Siddique KHM, Pandey MK, Farooq M, Varshney RK, Chapman MA, Boesch C, Daszkowska-Golec A, Foyer CH. Enhancing climate change resilience in agricultural crops. Curr Biol [Internet]. 2023 Dec;33(23):R1246–61. Available from: https://linkinghub.elsevier.com/retrieve/pii/S096098222301429X

100. Khawula S, Daniel AI, Nyawo N, Ndlazi K, Sibiya S, Ntshalintshali S, Nzuza G, Gokul A, Keyster M, Klein A, Niekerk L-A, Nkomo M. Optimizing plant resilience with growth-promoting Rhizobacteria under abiotic and biotic stress conditions. Plant Stress [Internet]. 2025 Sep;17(June):100949. Available from: 10.1016/j.stress.2025.100949

101. Pandey GR, Silwal S, Shrestha A, Pokharel S, Khanal BC, Acharya R. Defying Salinity, Drought, and pH Extremes: A Multifunctional Rhizobacterium, *Burkholderia gladioli* ST3M-39a, Matches Fertilizer Efficacy in Wheat via Phosphate Solubilization [Internet]. Mendeley Data; 2025. Available from: https://data.mendeley.com/datasets/h7g7cp3jxk

